# Limbic System White Matter in Children and Adolescents with ADHD: A Longitudinal Diffusion MRI Analysis

**DOI:** 10.1101/2024.09.17.613410

**Authors:** Michael Connaughton, Alexander Leemans, Timothy J. Silk, Vicki Anderson, Erik O’Hanlon, Robert Whelan, Jane McGrath

## Abstract

Attention-deficit/hyperactivity disorder (ADHD) is increasingly recognized as a disorder linked to atypical white matter development across large-scale brain networks. However, current research predominantly focuses on cortical networks, leaving the developmental trajectories of many subcortical networks, including the limbic system, largely unexplored. The limbic system is crucial for emotion and cognition, making it a key area of interest in ADHD research. This study employed multi-shell high angular resolution diffusion magnetic resonance imaging to map the development of limbic system white matter in individuals with ADHD (n = 72) and controls (n = 97) across three time points between ages 9 and 14. Diffusion kurtosis imaging and graph theory metrics were used to characterize limbic system white matter, alongside assessments of emotional regulation and ADHD symptom severity. Compared to controls, individuals with ADHD exhibited significantly lower microstructural organization, particularly in kurtosis anisotropy, within the bilateral cingulum bundle from childhood to adolescence. Brain-behavior analyses further revealed that higher ADHD symptom severity was associated with a lower number of limbic system white matter connections, notably decreased routing efficiency and network density. These findings offer novel insights into the role of disrupted limbic system white matter in ADHD pathophysiology, broadening our understanding of the disorder’s neural mechanisms and opening promising avenues for future exploration of subcortical brain networks.

## Introduction

Attention-deficit/hyperactivity disorder (ADHD) is increasingly viewed as a disorder of brain connectivity, specifically microstructural differences in the white matter tracts that interconnect large-scale brain systems (1, 2). White matter, also known as axons, is the tissue that connects various brain regions, and its development is crucial for supporting neurotypical brain function and cognition (3, 4). Healthy white matter development involves a highly conserved sequence of biological processes across species during early life that enable the formation, maintenance and optimization of the brain’s interconnections (5). Disruptions to these stages can have lasting cognitive, behavioral and neurological consequences (6). Two neurodevelopmental processes that appear particularly important in ADHD are neural formation and myelination (7). Neurotypical myelination involves the formation of the myelin sheath, which insulates axons and enhances the speed of signal transmission between neurons, thereby supporting optimal neural processing (7). Neural formation, specifically neuronal pathfinding, refers to the ability of neurons to establish correct connections during development, which is essential for effective communication between brain regions and is crucial for coordinated cognitive and behavioral functions (8). It has been proposed that the neural phenotype observed in ADHD may be caused by disruptions to these neurodevelopmental processes (7).

The transition from childhood to adolescence is a period marked by significant changes in ADHD symptom expression (9). Most individuals experience some improvement, particularly in hyperactivity-impulsivity, while symptoms of inattention tend to be more persistent (9). This period is also crucial for brain development, characterized by substantial white matter reorganization, including increased myelination and refined neural connections (3, 10). Therefore, it is possible that alterations in brain white matter development are linked to some of the variability in ADHD symptom trajectories observed during this critical developmental window.

Currently, the most common in-vivo measure of white matter is diffusion magnetic resonance imaging (MRI) (11). This imaging technique is optimized to capture the microstructural properties of white matter, providing indirect measures of the underlying neurobiological mechanisms of connectivity, organization, and integrity (11). Although sparse, current longitudinal diffusion MRI research has identified widespread developmental differences in individuals with ADHD compared to controls (12–14). These differences are characterized by ADHD-associated reductions in white matter organization in thalamic, frontostriatal, and corticospinal pathways (12, 13), as well as reductions in local white matter interconnectivity in the cingulate and middle cortex regions (14). These differences support the theory that ADHD may be linked to disrupted myelination processes during brain development (7). While longitudinal studies have provided valuable insights into the potential underlying mechanism of ADHD, much of the current diffusion MRI research has focused on cortical structures and networks, such as the frontostriatal, default mode, and ventral attention networks (15). This cortical focus is particularly surprising given that MRI research suggests structural abnormalities in subcortical brain regions may be a core feature of ADHD throughout a lifetime (16). To further understand the neurobiological mechanisms of ADHD, it is essential to investigate the development of subcortical structures and networks.

A promising – yet underexplored – brain network in ADHD pathophysiological research is the limbic system, a subcortical brain network that links visceral states with emotional cognition and behavior (17, 18). A focus on the limbic system is particularly relevant given the high prevalence of emotional dysregulation in ADHD, a symptom that can have significant long-term impacts on an individual’s social, academic, and psychological well-being (19). While atypical limbic system structure and function have been identified in various neurodevelopmental disorders (20), its role in ADHD remains largely unknown. Recently, using the same longitudinal MRI cohort as this study, we discovered that atypical grey matter development in the limbic system may be a key neurobiological characteristic of ADHD (21). Specifically, individuals with ADHD displayed fixed, non-progressive volumetric reductions bilaterally in the amygdala, hippocampus, orbitofrontal cortex and cingulate gyrus, compared to non-ADHD controls, during the transition from childhood into adolescence (21). Given the growing evidence of disrupted brain connectivity as a core feature of ADHD (1, 2), investigating the maturation of limbic system white matter during a developmentally sensitive period—when significant shifts in ADHD symptoms commonly occur—is a crucial next step that could provide valuable insights into the neurobiological underpinnings of the disorder.

Historically, exploring complex subcortical structures such as limbic system white matter tracts has been challenging for diffusion MRI researchers. Since the early 2000s, the most common diffusion MRI modelling technique has been Diffusion Tensor Imaging (DTI) (22). While a pioneering technique, DTI’s inability to model crossing fibers has presented a major limitation for white matter reconstruction (22). However, advancements in diffusion MRI acquisition techniques, such as multi-shell High Angular Resolution Diffusion Imaging (HARDI), have enabled fiber tracking techniques, such as constrained spherical deconvolution (CSD) (23) that can overcome many of the limitations of the DTI model (24). These improved fiber tracking methods enhance reconstruction accuracy and, importantly, can model crossing fibers which is essential for reliably reconstructing complex subcortical pathways (25). The increased diffusion profile data provided by multi-shell HARDI can then be modelled using higher-order techniques such as Diffusion Kurtosis Imaging (DKI) (26), an advanced MRI technique that provides a more detailed picture of brain tissue by measuring how water molecules move in complex cellular environments (27). Unlike traditional DTI, which assumes water diffusion follows a simple Gaussian distribution, DKI captures non-Gaussian diffusion patterns, offering a more accurate representation of the brain’s microstructure (27). DKI captures more detailed information about the diffusion profile than DTI, making it more sensitive to neurobiological changes, including demyelination—a proposed feature of ADHD (7, 22, 28, 29). A DKI metric that has demonstrated the ability to detect myelination is kurtosis anisotropy (KA) (30, 31). KA measures the deviation of water diffusion from a simple pattern, as expected in healthy axons (30, 31). Higher KA values indicate healthier, more organized white matter with intact myelin, while lower values suggest demyelination and damage (30, 31). Details of other DKI metrics and their links to neurobiological processes are provided in Table 1.

**Table 1.**
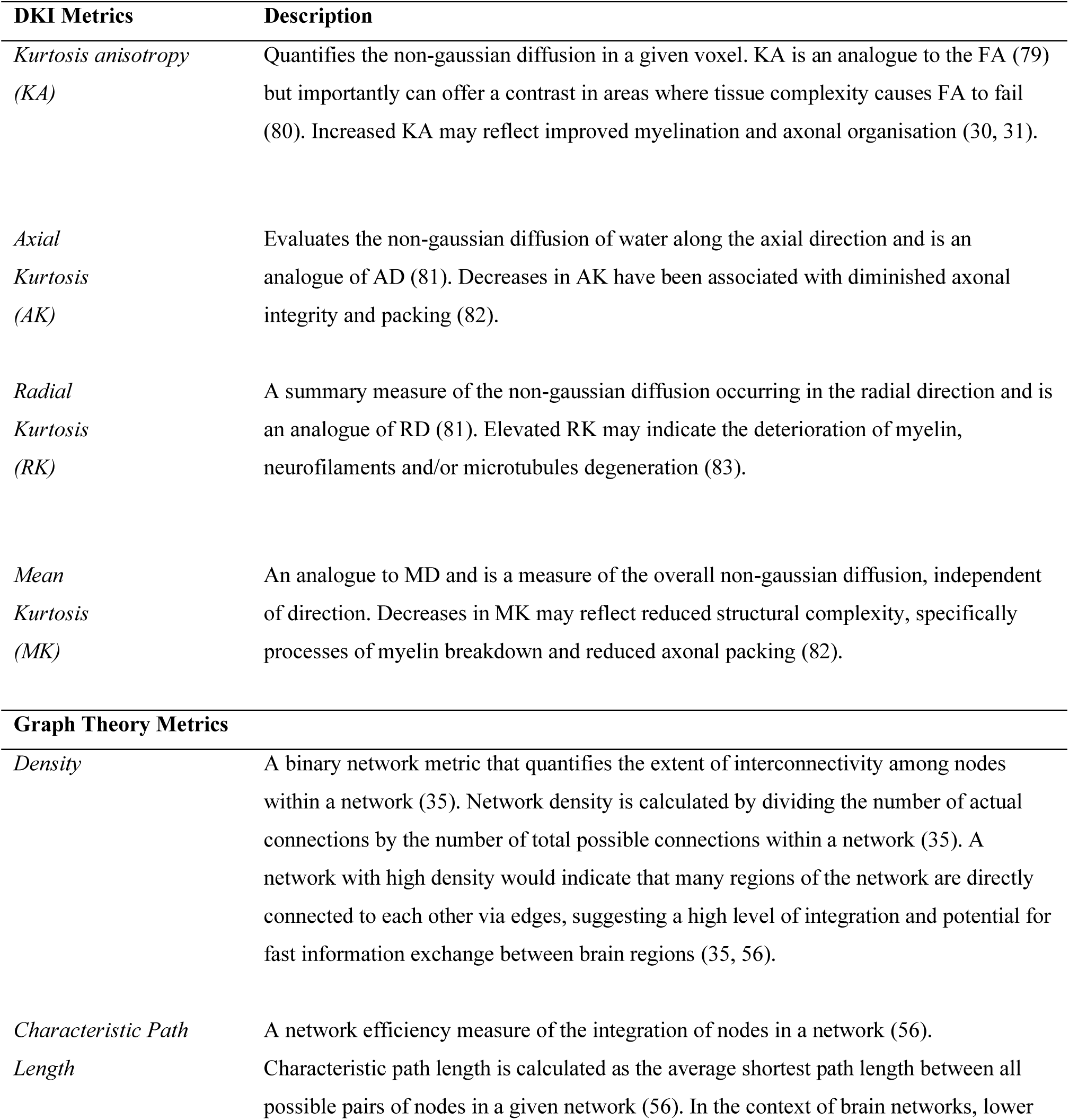
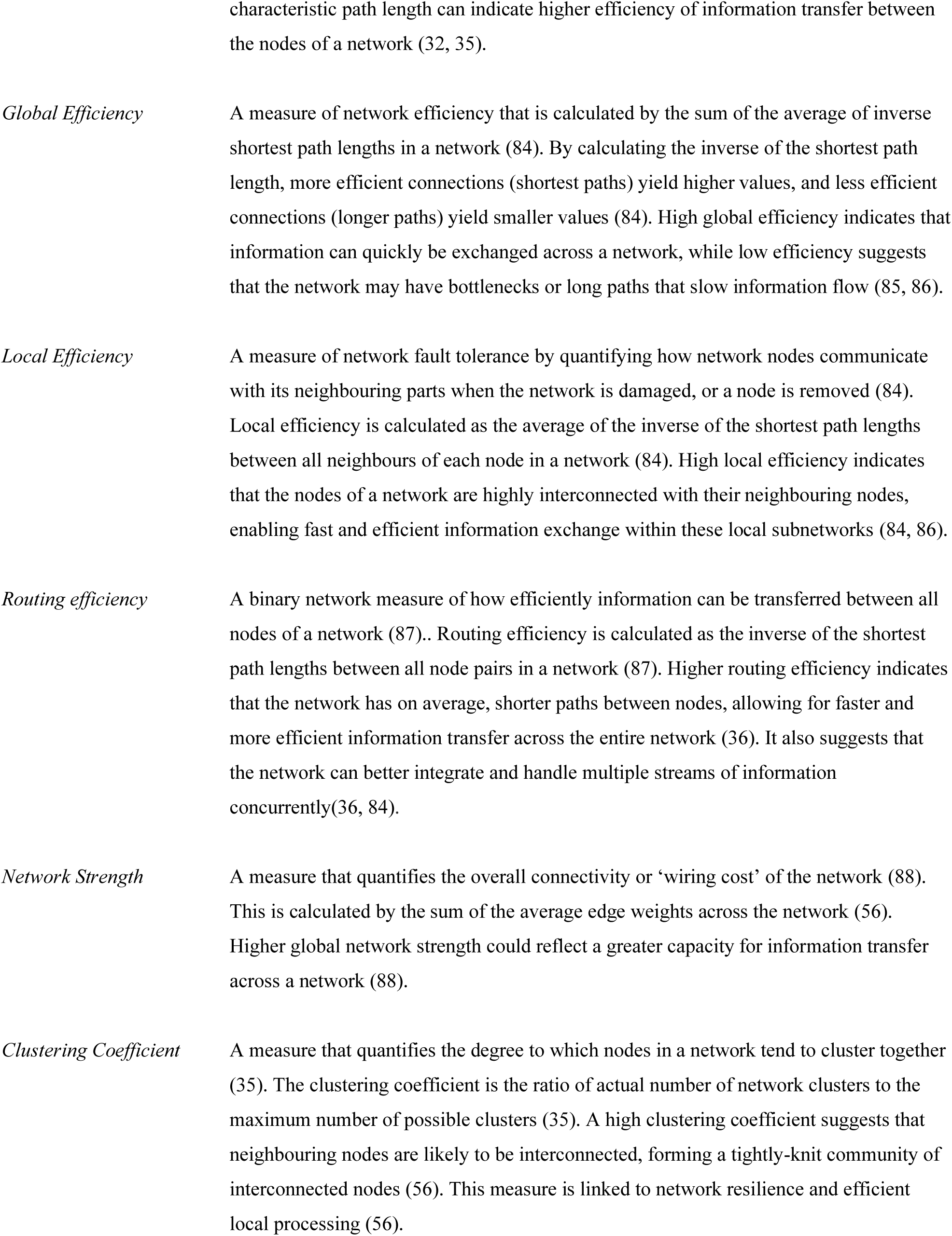
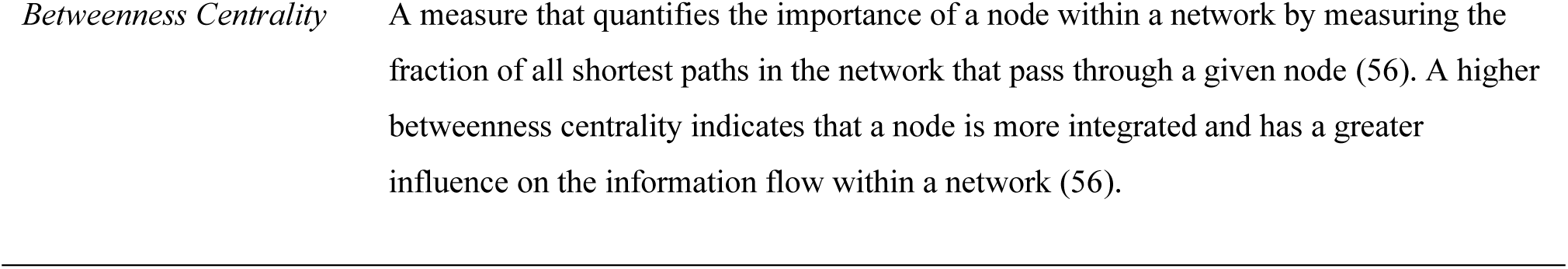
Description and clinical implications of micro – and macrostructural diffusion MRI metrics.

In addition to microstructural analysis, macrostructural white matter network analyses can provide valuable information on the topology of brain white matter, detailing the degree of interconnectivity across brain structures. In white matter network analysis, a structural connectivity matrix is constructed by modelling the brain as a network of nodes (grey matter regions generated by structural MRI) and their interconnecting edges (white matter fibers generated from diffusion tractography) (32). This matrix can then be characterized using *graph theory*, a branch of mathematics that offers a powerful framework for quantifying the overall organization and efficiency of information transfer across brain networks (33). Through this framework, a variety of graph-theoretical metrics can be derived to provide information about brain network properties, such as efficiency, centrality, modularity, and clustering, offering a comprehensive understanding of how information is processed and integrated across different brain regions via white matter tracts (32) (Table 1). Additionally, these metrics have been shown to serve as proxy measures for neurobiological processes underlying early life white matter development, including neural connection formation and synaptic pruning (34). Binary network metrics are particularly useful as they construct networks where each connection between brain regions is either ‘present’ (assigned a value of 1) or ‘absent’ (assigned a value of 0). This method allows researchers to analyze structural connectivity by mapping the trajectories of white matter pathways and identifying whether specific connections have formed. Two common binary metrics are network density and routing efficiency, which reflect the number of interconnected nodes in a network; lower values in these metrics indicate decreased white matter interconnectivity of the network (35, 36). Given the evidence that ADHD is associated with disruptions in neural formation during early brain development (7), this study seeks to determine if similar disruptions can be observed in the white matter of the limbic system.

Using three timepoint multi-shell HARDI data collected during the transition from childhood to adolescence, this study aimed to 1) investigate the differences in limbic system white matter development between individuals with ADHD and non-ADHD controls, and 2) explore the relationship between ADHD symptom severity and limbic system white matter development. Based on previous findings (1, 7, 21), it was hypothesized that individuals with ADHD would exhibit 1) lower kurtosis anisotropy and 2) lower white matter interconnectivity of the limbic system compared to non-ADHD controls. Additionally, we expected that greater severity of ADHD symptoms, including emotional dysregulation, would be associated with 1) lower microstructural organization and 2) lower interconnectivity of limbic system white matter during this developmental period.

## Methods and Materials

### Study Design and Participants

This longitudinal study involved 169 children (72 with ADHD and 97 controls) recruited for the Neuroimaging of the Children’s Attention Project (NICAP) (37). NICAP is a multimodal longitudinal neuroimaging study conducted at a single site, examining a community-based cohort of children with and without ADHD at three intervals over five years, from ages 9 to 14. Children in the ADHD group were diagnosed with ADHD using the National Institute of Mental Health Diagnostic Interview Schedule for Children (DISC-IV) (38) at each assessment point (recruitment, wave 1, and wave 3 imaging time points). The control group participants did not meet the diagnostic criteria for ADHD at any assessment point. Written informed consent was obtained from the parents or guardians of all participants prior to enrolment. The study received ethical approval from both the Royal Children’s Hospital Melbourne Human Research Ethics Committee (HREC #34071) and the School of Psychology, Trinity College Dublin (SPRECC042021-01).

### Assessment of Clinical and Emotional Metrics

At each of the three NICAP study intervals, the Conner’s 3 ADHD Index (CAI), reported by parents, was used to evaluate the core symptoms of ADHD in participants, where higher scores indicated a greater presence of symptoms. (39). The Affective Reactivity Index (ARI) was administered at the first and second time points to assess emotional reactivity and irritability, with higher scores indicating increased emotional dysregulation and irritability (40). Matrix reasoning scores (based on Wechsler Abbreviated Scale of Intelligence (41) – mean matrix reasoning T-score), whereby higher scores indicate better reasoning ability, were collected at all three time points. Socioeconomic status (SES) scores (determined using the Socio-economic Indexes for Areas (42)) were collected at all three time points, where higher scores indicated higher SES.

### MRI Data Acquisition

All neuroimaging data were collected at a single site at the Murdoch Children’s Research Institute at the Royal Children’s Hospital, Melbourne, on a 3-Tesla Siemens scanner using a 32-channel head coil. However while the first two waves were collected on a total imaging matrix (TIM) Trio scanner, the third wave was collected after an upgrade to a MAGNETOM Prisma scanner (previous papers on this cohort have found minimal effects of scanner upgrade (43)). Diffusion MRI data were collected using a multi-band accelerated EPI sequence protocol (CMRR, University of Minnesota) (44), which allows for accelerated diffusion MRI multi-shell acquisition. Multi-shell high angular resolution diffusion imaging (HARDI) data were acquired using the following protocol: 2800 s/mm^2^, (60 directions, four interleaved b = 0 volumes), 2000 s/mm^2^ (45 directions, 6 interleaved b = 0 volumes), and 1000 s/mm^2^ (25 directions, 6 interleaved b = 0 volumes) with an anterior-posterior phase encoding direction.

### MRI Data Processing

#### Diffusion MRI

Diffusion images were processed using ExploreDTI software (45). The processing procedures used in this study followed a comprehensive diffusion MRI processing step-by-step guide (46). Briefly the pre-processing steps included signal drift, Gibbs ringing, echo-planar imaging (EPI) distortion and subject motion correction. To reduce the effects of EPI distortion and subject motion, each diffusion MRI scan was co-registered to a subject-specific structural MRI T1 image (Section 1 in supplemental material for T1 image processing details). As recommended, b-matrices were rotated while correcting for subject motion/EPI distortion (47). This ensures that the diffusion gradients remain accurately aligned with the brain’s anatomy to improve the integrity of the diffusion metrics (48). Robust tensor estimation through the REKINDLE method was selected for its efficacy in removing outliers and maximizing tensor accuracy, which is particularly beneficial in white matter fibres vulnerable to corticospinal fluid contamination, such as those in the limbic system (49, 50).

This study used deterministic Constrained Spherical Deconvolution (CSD) tractography to reconstruct whole brain white matter fibres, using the recursive calibration of the response function (23). Whole-brain fibre deterministic tractography was performed using the following settings: seed points resolution 2x2x2 mm^3^, step size 1 mm, angle threshold 60°, fibre length range, 10 – 500 mm. Quality control procedures used in this study are described in Section 2 of the supplemental material (eFigure 1). Following quality control procedures, 15 scans were excluded from the DKI analysis and 22 scans were excluded from the connectomics analysis.

**Figure 1.**
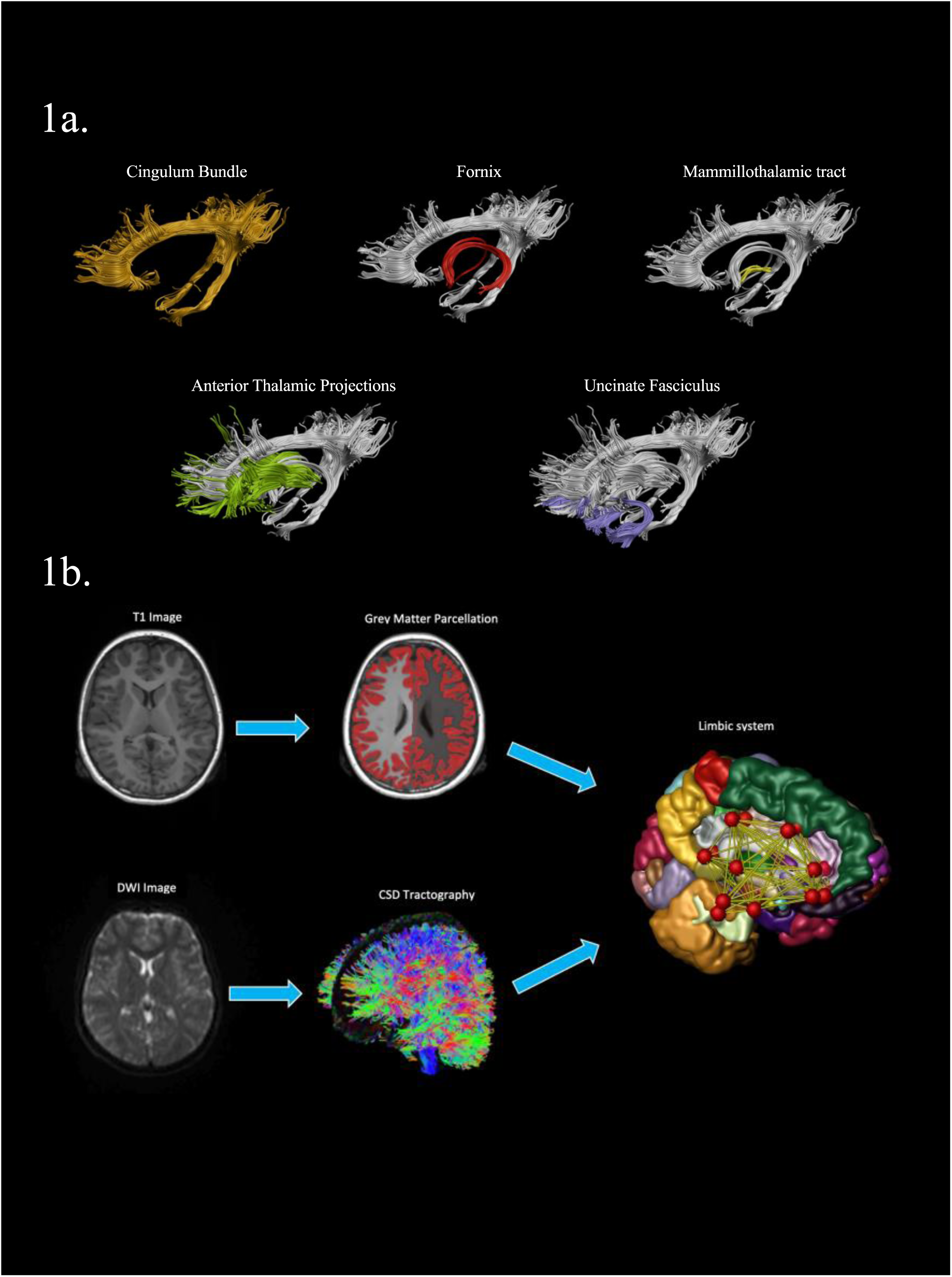
Visual guide to limbic system white matter extraction procedures used in this study. **a) r**econstructed limbic system white matter tracts following manual ROI tractography, each fibre were extracted bilaterally for each participant in this study. **b)** This figure outlines the key processing steps for the construction of a subject-specific limbic system connectome. After processing T1 and diffusion MRI images, full cortical parcellation and whole-brain tractography is performed. During processing the diffusion MRI image was co-registered to the T1 image, for connectomics construction it is essential that these two images are registered to the same space. Using ExploreDTI software the nodes are defined and edges are then generated using the “pass” definition, which assigns a streamline to a pair of nodes if the streamline intersects them. These procedures are repeated for each subject in the study to create subject-specific limbic system connectomes.

#### Manual Tractography for Limbic System White Matter

To ensure the highest accuracy and reliability in limbic system white matter reconstruction, this study employed manual region of interest (ROI) tractography (51). Despite being time-intensive, manual methods provide the highest inter-rater and intra-rater reliability compared to automated techniques due to their incorporation of expert neuroanatomical knowledge and subject-specific approach (51). The five major white matter tracts of the limbic system (cingulum bundle, fornix, mammillothalamic tract, anterior thalamic projections, and uncinate fasciculus) were isolated bilaterally for all participants in this study (Figure 1). Full details of tract extraction protocols are provided in Section 3 of the supplemental material. Once reconstructed, the microstructural organisation of these white matter tracts were quantified using DKI metrics, specifically kurtosis anisotropy (KA), mean kurtosis (MK), radial kurtosis (RK), and axial kurtosis (AK) (Table 1).

#### Limbic System White Matter Connectome Construction

ExploreDTI software was used to construct subject-specific networks, with nodes and edges as the fundamental elements (52). Nodes were defined using automated segmentation and parcellation from the Destrieux Atlas, based on subject-specific T1 MRI images (53, 54). Bilateral anatomical masks were created for the following structures: thalamus, hippocampus, amygdala, rostral anterior cingulate gyrus, caudal anterior cingulate gyrus, posterior cingulate gyrus, isthmus cingulate gyrus, parahippocampal gyrus, lateral orbitofrontal cortex, and medial orbitofrontal cortex. Using whole-brain CSD tractography, edges were defined with Hagmann weighting, where the streamline count reflects structural connectivity (55). Streamlines were assigned to node pairs if they intersected them in their path without further restrictions on their start or endpoint (i.e., the network was unrestricted, allowing potential connections between every pair of nodes). A visual guide is provided in Figure 1.

Using the Brain Connectivity Toolkit (56), graph theory metrics were used to measure the macrostructural organisation of limbic system white matter. The global graph theory metrics used in this study were global efficiency, characteristic path length, network density, clustering coefficient, network strength, local efficiency and routing efficiency (Table 1).

#### Statistical Analysis

Before commencing statistical analyses a systematic inspection of the data including outlier identification was performed. Outliers were defined as any data points diverging by ±3 standard deviations from the data mean. In line with established statistical practice, these identified outliers were removed from the statistical analysis (57).

Full details and breakdown of the statistical analyses used in this study are provided in Section 4 of the supplemental material. Briefly, to measure between-group differences in volume of limbic system structures, mixed-effects modelling was performed using the *lme4* package in R (version 1.1-27.1) (58). Sex, age, age at baseline and intracranial volume (for graph theory analysis) were included as covariates. Given prior work suggesting non-linear relationships between brain structure and age (59), both linear and quadratic models were tested to identify the optimal growth curve for each structure. The details of the mixed models tested in this study are presented in eTable 1. Brain-behaviour statistical analyses were conducted to examine the relationship between microstructural organization of limbic system white matter tracts and behavioural measures (Conner’s 3 ADHD Index and Affective Reactivity Index) in children and adolescents with ADHD. The models used for brain-behavior analyses are presented in eTable 1.

A post hoc power analysis was conducted to evaluate the statistical power of the study in detecting non-significant findings. The analyses investigated small (Cohen’s d = 0.20), medium (Cohen’s d = 0.30), and large (Cohen’s d = 0.40) effect sizes using the ‘powerlmm’ package in R (60), based on a longitudinal mixed-effects model. The models incorporated an intraclass correlation coefficient (ICC) of 0.5 and a stringent alpha level of 0.005 to account for multiple comparisons. A dropout rate of 28% was modeled using a Weibull distribution, reflecting the observed dropout rate in the study.

Multiple comparisons were handled using a two-stage false discovery rate (FDR) correction, using the *MuToss* package (61) in R (v.4.1.1). Additional sensitivity analyses were conducted to evaluate the potential influence of confounding factors, such as case-control sex imbalance, ADHD medication status, comorbidity status, and head motion on the statistical analyses.

## Results

### Demographics and Clinical Characteristics of Study Population

Following MRI quality control, the final sample comprised of 360 diffusion MRI scans (154 ADHD, 206 controls) for the DKI analyses and 342 scans (149 ADHD, 193 controls) for the connectomics analyses. The comprehensive demographic and clinical characteristics of the study population can be found in Table 2. There were no significant differences between the groups in terms of age, handedness, socioeconomic status, or matrix reasoning at any of the three study time points. Children diagnosed with ADHD exhibited significantly higher ADHD symptom severity compared to the control group, as expected. Additionally, children with ADHD had a higher presence of co-morbid externalizing and internalizing disorders compare to controls. At any given wave 22.5% – 30.5% of the ADHD group were taking medication related to their diagnosis (eTable 2 for full medication use breakdown).

**Table 2.**
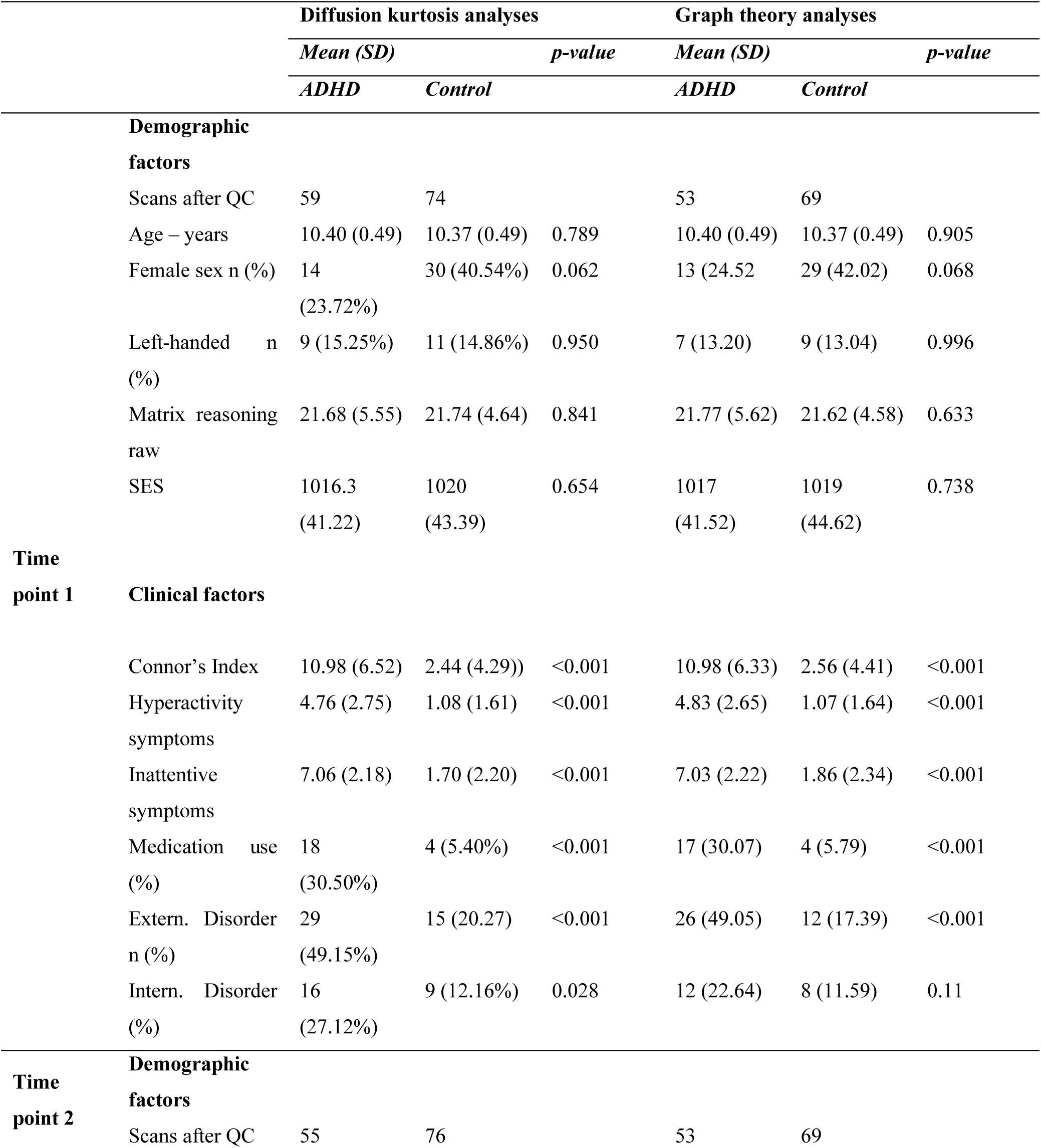
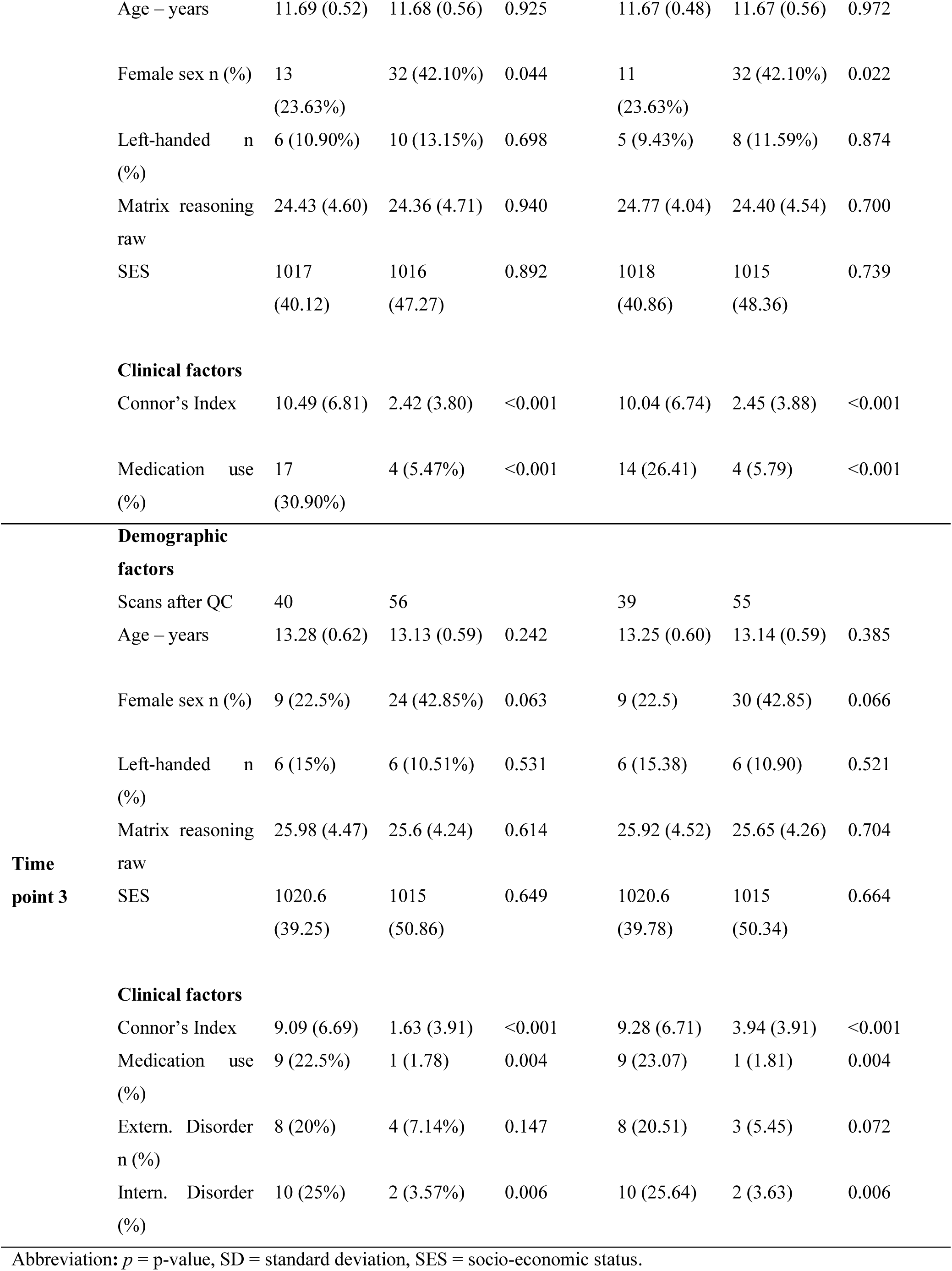
Demographics and clinical variables of study participants.

### Limbic System White Matter in Children and Adolescents with ADHD and Controls

Full details of the top-down mixed model selection analyses are provided in eTable 3-6. Using the optimally selected models, the results found that individuals with ADHD had significantly lower kurtosis anisotropy in the bilateral cingulum bundle across all three study time points compared to controls, with results surviving post-hoc analysis (Left, β = -0.36, 95% CI = -0.63 to -0.09; Right, β = -0.36, 95% CI = -0.62 to -0.11, (Figure 2)). There were no other significant group differences in limbic system white matter DKI or graph theory metrics among individuals with ADHD and controls. There was no significant difference in the group-by-age interaction regarding the limbic system white matter metrics among individuals with ADHD and controls. This finding suggests that the developmental trajectories of limbic system white matter did not significantly differ between the ADHD and control groups. All between-group analyses are presented in eTables 7-10 and eFigure 2.

**Figure 2.**
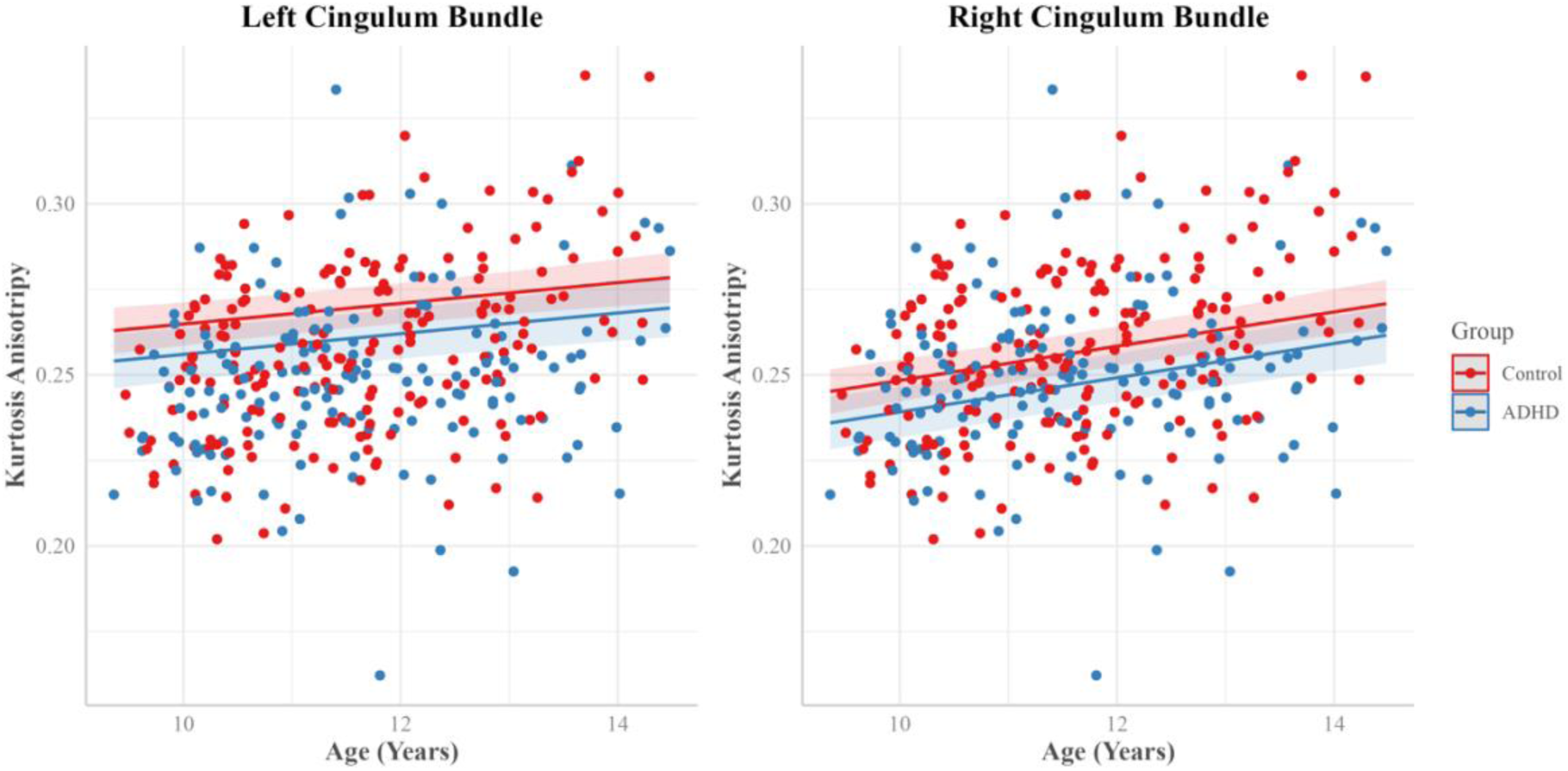
Kurtosis anisotropy of the bilateral cingulum bundle across the three study time points. Bilateral cingulum bundle KA in the ADHD and control groups during the transition from childhood to mid-adolescence. A main effect of group interaction was observed in the bilateral cingulum; the ADHD group displayed reduced KA compared to the control group across all three study time points.

### ADHD Symptoms Severity and Limbic System White Matter in ADHD

The brain-behavior analysis found a significant effect of Conner’s 3 ADHD index scores on routing efficiency (β = - 0.30, 95% CI = -0.49 to -0.10) and network density (β = - 0.25, 95% CI = -0.44 to -0.06) in the ADHD group across the three study time points, with results surviving post-hoc analysis (Figure 3). This suggests that decreased routing efficiency and network density in the limbic system were significant predictors of heightened ADHD symptom severity among individuals with ADHD throughout childhood and adolescence. There was no other significant effect of CAI or ARI on any other limbic system white matter metric in children and adolescents with ADHD. All brain-behaviour findings are presented in eTables 11-14 and eFigures 3-6.

**Figure 3.**
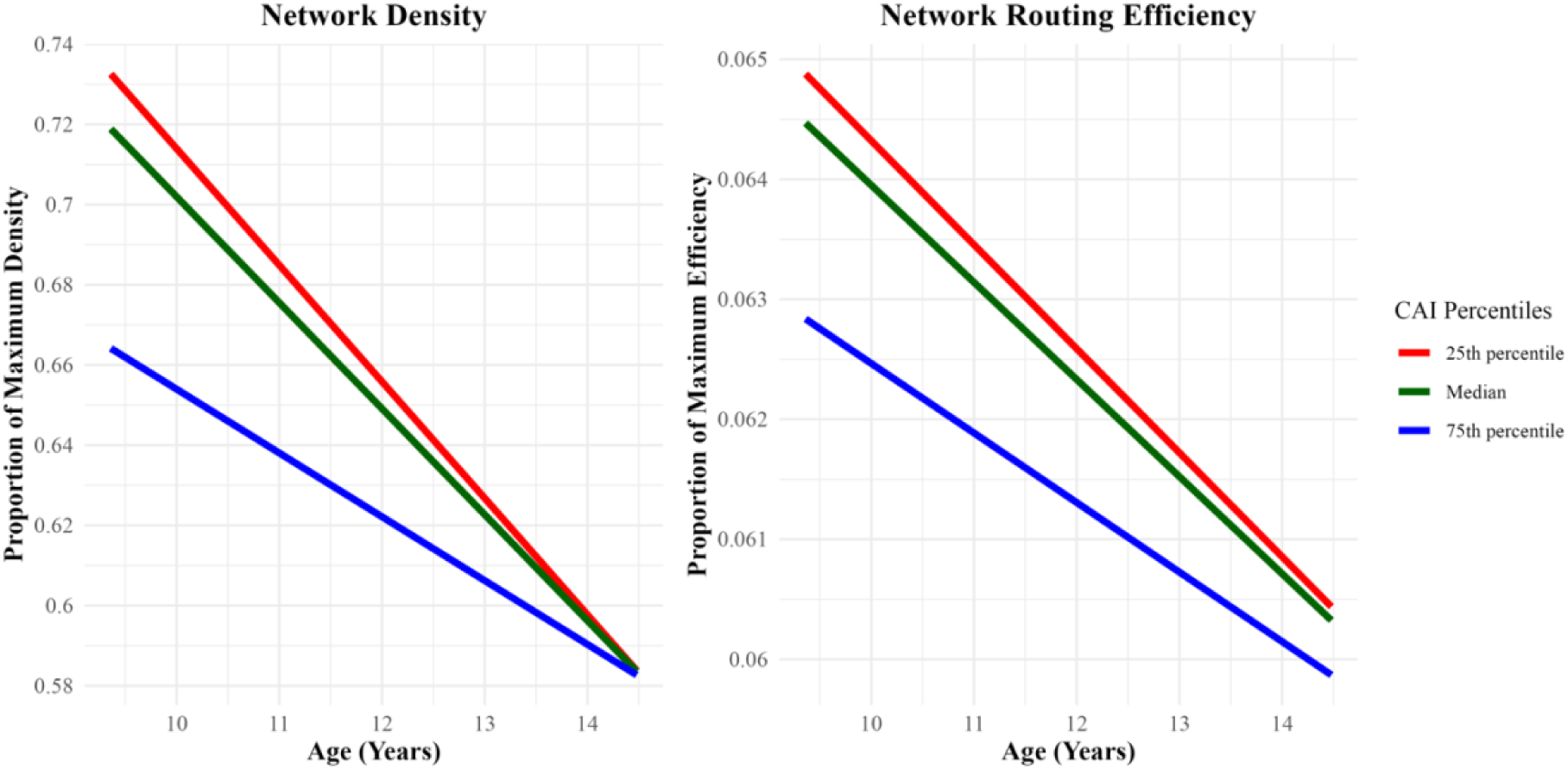
Relationship between of CAI score and graph theory metric in the ADHD group over the NICAP study time points. This figure illustrates the predicted change in limbic system network measure (y-axis) as a function of age (x-axis). The colours represent different percentiles of CAI scores: 25^th^ percentile (red), median (green), and 75^th^ (blue). The Y axis represents the predicted routing efficiency values from the LMM model used in the brain- behaviour analysis.

### Power Analyses

The power analysis for these parameters indicated a power of 1.00 (SE = 0.0144) for detecting effect sizes of 0.3 and 0.4, and a power of 0.91 (SE = 0.0140) for detecting an effect size of 0.2. These results demonstrate that the study was highly sensitive to medium and large effects and adequately powered to detect small effects. Consequently, the non-significant findings are likely not due to insufficient power but rather suggest the absence of substantial effects within the study parameters.

### Sensitivity Analyses

The reported statistically significant results survived all sensitivity analyses, demonstrating that the results were most likely not biased by the sex ratio per group, medication, comorbidity status or head motion. Full details of sensitivity analyses are presented in the supplemental material (eTables 15-21 and eFigure 9-12).

## Discussion

The objectives of this longitudinal study were to: 1) examine the development of limbic system white matter in individuals with ADHD compared to non-ADHD controls, and 2) explore the relationship between ADHD symptom severity and limbic system white matter development during the transition from childhood to adolescence. As hypothesized, individuals with ADHD exhibited lower microstructural organization in the white matter bundles of the limbic system, specifically in the bilateral cingulum bundles, compared to non-ADHD controls during this developmental period. Furthermore, brain-behaviour analyses revealed that lower interconnectivity of limbic system white matter was linked to greater symptom severity in individuals with ADHD. Two hypotheses were not supported. No between-group difference was observed in the topological organization of limbic system white matter. Additionally, no significant association was found between limbic system microstructural organization and ADHD symptom severity. Importantly, post hoc power analysis confirmed that the study was adequately powered to detect meaningful effects, reinforcing that the non-significant findings genuinely reflect the absence of substantial effects within the study parameters.

This discussion examines the findings of the current study in relation to other pathophysiological ADHD research, focusing on two key areas, 1) the development of the cingulum bundle white matter and its implications for ADHD pathophysiology, and 2) the relationship between limbic system white matter connectivity and ADHD symptom severity.

### Cingulum Bundle Development and ADHD Pathophysiology

This study found that individuals with ADHD displayed lower microstructural organization, specifically lower kurtosis anisotropy (KA; indicating demyelination and damage), in the cingulum bundle bilaterally compared to controls across childhood and into mid-adolescence (Figure 2). In both groups, cingulum white matter organization increased linearly across the three NICAP study time points, a pattern consistent with neurotypical cingulum bundle maturation (62, 63). This suggests that in ADHD, although the developmental trajectory of cingulum bundle white matter microstructure follows a neurotypical pattern, it is characterized by a fixed, non-progressive reduction in microstructural organization across childhood and mid-adolescence. The cingulum bundle integrates cognitive and emotional processes through its connections with the frontal, parietal, temporal, and subcortical regions (64). Altered microstructural organization of the cingulum bundle has been linked to a wide range of functional deficits, including issues with executive function, attention, emotional processing, and regulation (65). Thus, cingulum bundle developmental alterations may contribute significantly to the cognitive and emotional challenges observed in individuals with ADHD, underscoring the importance of this neural pathway in the disorder’s pathology.

The results of this study are mirrored in another MRI study on the same NICAP data set (21). Specifically, a structural MRI study, focusing on grey matter, showed that during childhood and mid-adolescence, ADHD-associated fixed, non-progressive volumetric reductions were also observed in the interconnected limbic system regions of the cingulum bundle, specifically the cingulate gyrus, orbitofrontal cortex, amygdala, and hippocampus (21). This suggests that the cingulum bundle and its interconnections may form a subnetwork within the limbic system that could be central to the disrupted neurodevelopment seen in ADHD.

Neurobiologically, lower KA has been shown to be a sensitive marker of demyelination (30, 31). As such, the findings of this study support the hypothesis that the neurobiological phenotype seen in ADHD may be underpinned by dysregulated myelination, particularly in the frontostriatal and limbic circuits (7, 28). A recent genome-wide association study (GWAS) identified the most significant ADHD-associated locus as the gene encoding the beta-galactoside-alpha-2, 3-sialyltransferase-III (ST3GAL3) membrane protein (66). ST3GAL3 plays a key role in the sialylation of glycoproteins, a process essential for the proper formation and function of the myelin sheath (67). Animal research has found that ST3GAL3-deficient mice display ADHD-like behaviours (67). Additionally, these ADHD behavioural phenotypes were driven by altered myelination, characterized by a reduction in major myelin proteins, fewer myelinated axons, and an overall decrease in myelin (67). As such, ADHD-associated developmental disruptions in myelination processes may underpin the fixed, non-progressive reduction in cingulum bundle KA seen in this study.

### Limbic System Connectivity and ADHD Symptom Severity

This is the first study to investigate the relationship between the development of limbic system white matter interconnectivity and ADHD symptom expression during the transition from childhood to early adolescence, a period characterized by significant symptom fluctuation (19). The findings revealed that lower limbic system network density and routing efficiency— both measures of network interconnectivity (35, 36)—were significantly linked to greater ADHD symptom severity from childhood into early adolescence (Figure 3). This suggests that during this transition, limbic system regions are less interconnected in individuals with greater ADHD symptoms compared to those with milder symptoms. As optimal brain function relies on the efficient transfer of information between brain regions via white matter tracts (68), this reduced interconnectivity may disrupt limbic system functioning. Disrupted communication and coordination between regions involved in emotion regulation, attention, and impulse control may exacerbate core ADHD symptoms such as inattention and impulsivity (17). In contrast to severe ADHD cases, milder ADHD cases displayed less deviation from non-ADHD controls. For instance, individuals with average ADHD symptom severity had a mean density of 0.630 and a mean routing efficiency of 0.0614. In comparison, non-ADHD controls exhibited a mean density of 0.639 and a mean routing efficiency of 0.0618. However, individuals with high ADHD symptom severity (1 SD above the mean CAI score) showed more pronounced deviations, with a mean density of 0.620 and a mean routing efficiency of 0.0608. These data demonstrated that while average ADHD cases do show slight deviations in limbic system interconnectivity metrics, these deviations are less pronounced than those seen in high ADHD symptom severity cases. Therefore, among individuals with milder ADHD, while their interconnectivity may not be optimal, they could still remain within a functional range, allowing for better overall function and fewer symptoms. This may explain why within-group analysis showed an association between limbic system interconnectivity and symptom severity, yet no significant difference was found between ADHD and non-ADHD controls. Thus, limbic system interconnectivity may be a sensitive marker specifically for severe ADHD cases.

During adolescence, a neurodevelopmental process called "frontalization" occurs, where white matter connections to frontal regions are strengthened while those to subcortical regions are weakened (69). This shift reduces the autonomy of subcortical regions, making them more receptive to top-down control, which coincides with increased behavioural regulation during the transition to adulthood (69). The study observed this pattern of subcortical white matter refinement. However, individuals with severe ADHD displayed lower limbic system interconnectivity from baseline (around age 9) compared to those with milder cases, suggesting that individuals with more severe ADHD symptoms may have experienced early life disruptions in white matter formation. This observation aligns with other pathophysiological research indicating that ADHD pathophysiology may be connected to the disruption of neuronal formation processes during early brain development (7). Specifically, findings from large genome-wide association studies suggest that variants in ADHD-associated genes, such as CDH13 and PCDH7, might disrupt the normal formation of white matter fibres in the brain (66, 70, 71). This possible disruption in early brain neural formation processes could potentially explain the observed association between ADHD symptom expression and lower white matter interconnectivity seen in this study. While promising, it is important to note that our current understanding of neural formation in the context of ADHD is still limited. Continued longitudinal diffusion MRI research into underexplored brain regions and networks can provide vital insights into the neurodevelopmental trajectories underlying ADHD. These insights, when leveraged with current neurobiological theories, may help elucidate the mechanisms of the disorder.

### Limitations

This study had some limitations. For instance, while the study used a scientifically defined limbic system (17), it must be acknowledged that there is currently no universally accepted anatomical definition of the limbic system (72). Definitions vary, with some incorporating structures like the septal nucleus, insula, and parahippocampal gyrus (73). This variability in structural definitions presents a challenge for limbic system research, hindering the formation of a unified body of literature and a coherent understanding of limbic system functions.

Reliance on the Affective Reactivity Index (ARI) to measure emotion regulation, which may have contributed to the lack of observed association between emotion dysregulation and limbic system white matter. Although the ARI is commonly used to measure emotional dysregulation, it may not fully capture this multifaceted construct, as it primarily measures irritability and reactivity, which are only components of emotional dysregulation (74). Considering the complex and multi-dimensional nature of emotional regulation, future research on children with ADHD would benefit from incorporating a more comprehensive range of assessments, such as the Emotion Regulation Checklist (75), and Temperament in Middle Childhood Questionnaire (76).

A further limitation is the lack of understanding of how puberty affects white matter development in relation to ADHD. The influence of puberty on ADHD remains uncertain. Given that the presentation of ADHD symptoms and neuropsychological functioning appear to change as an individual enters puberty (77), and considering white matter has been shown to be particularly sensitive to remodelling with exposure to pubertal hormones (78). Clinically, understanding the impact of pubertal hormones on brain development in ADHD could offer crucial insights into the progression and variability of the disorder during adolescence. This knowledge would allow for more accurate predictions of its clinical course and support the development of targeted interventions tailored to the unique challenges faced during this critical period.

## Conclusion

This study utilized advanced diffusion MRI techniques to provide novel insights into limbic system white matter development in a large cohort of children and adolescents with ADHD. The findings reveal how disruptions in this critical brain network are intricately linked to the disorder, offering strong support for leading pathophysiological theories of ADHD (7, 28). These results open up promising avenues for further research into the pathophysiology of ADHD, underscoring the importance of continued exploration of subcortical brain network

