## Supplemental Material for "Limbic System White Matter in Children and Adolescents with ADHD: A Longitudinal Diffusion MRI Analysis"

*Connaughton et al.,*

#### 1. Structural MRI Data Acquisition and Processing

T1-weighted volumes were collected using a multi-echo magnetization prepared rapid gradient-echo (MEMPRAGE) sequence with in-scanner motion correction (MoCo) (TR = 2530 ms, TE = 1.77, 3.51, 5.32, 7.2 ms, matrix = 256 x 232, number of slices = 176, voxel size = 0.9 $mm^3$, flip angle = $7^o$). T2-weighted volumes were obtained using a T2-SPACE (Sampling perfection with application optimized contrast with flip angle evolution) protocol (TR = 3200 ms, TE = 532, matrix = 256 x 230, slices = 176, voxel size = 0.9 $mm^3$). T1 and T2-weighted volumes were used to provide optimal sensitivity and increase the accuracy of subcortical brain reconstruction (Seiger et al. 2021; Iglesias et al. 2015).

FreeSurfer software (http://surfer.nmr.mgh.harvard.edu/) was used to isolate the structures of the limbic system. FreeSurfer analyses were performed using a Redhat-based scientific Linux 7 on the high-performance computing system at Trinity College Dublin, Ireland. All MRI images were processed using FreeSurfer's *recon -all* function (version 7.2) for full cortical reconstruction and brain segmentation (1, 2). The Desikan-Killiany-Tourville (DKT) atlas was used for brain parcellations (3).

#### 2. MRI Quality Control Procedure

MRI quality control (QC) procedures were undertaken at pre, during and post scan. Prior to the scan, a mock scanner session was completed to ensure participants were comfortable in the MRI environment and could minimise motion during live scanning session. During live scanning, Siemens in-scanner motion correction feature, adjusting the field-of-view and slice positioning in real-time to account for movement during the acquisition process (4). This feature significantly reduces the effect of motion artefacts and substantially improves image quality (5). This was particularly crucial for the studied population, namely with attentional and hyperactivity difficulties, where motion is a large challenge (6). In cases where head motion was high during scan acquisition, multiple scans were acquired until a suitable image was completed. Poor images due to motion were identified by the expert on-site radiographer.

#### 2.1. Tractography Quantality Control

Quality control of both raw and processed dMRI scans was conducted in line with diffusion MRI quality control guidelines (7). The visual inspection focused on identifying and correcting specific diffusion artifacts, including eddy currents, Gibbs ringing, chemical shifts, Nyquist ghosting, pulsations, interslice instabilities, and signal dropouts. In evaluating raw dMRI images, a 4-point Likert scale was utilized. A score of '1' was assigned to images free of visible artifacts, representing optimal quality. Images with a score of '2' exhibited minor artifacts, such as slight motion blur or minimal eddy current distortions, while a score of '3' was given to images with moderate artifacts, including more pronounced motion blur or eddy current distortions. Images severely compromised by artifacts, indicated by a score of '4', were characterised by excessive motion blur or significant susceptibility-induced geometric distortions. Any images that received a score of 3 or greater were excluded from the study. For the post-processed images, a 3-point Likert scale was employed to assess the effectiveness of artifact corrections and the accuracy of diffusion profiles. Images with near-perfect reconstructions were scored as '1', those with minor issues confined to small brain areas received a '2', and scans with poor reconstruction, evidenced by extensive or distorted areas, were scored '3' and excluded from the study. The study's methodology encompassed standardised viewing conditions, ensuring consistent lighting and display settings for image review. An experience neuroscientist, skilled in identifying and rating MRI artifacts, conducted the inspections. A sequential analysis approach was adopted; each raw image was initially reviewed for quality, followed by the assessment of processed images, guaranteeing an unbiased evaluation.

Following this thorough visual inspection, 360 scans were found suitable for analysis, with 15 scans excluded due to incomplete MRI image sets (missing b-value images) and 5 scans excluded due to quality concerns—2 during pre-processing and 3 post-processing. No manual edits were made to the data of the remaining scans.

#### 2.2. Connectomics Quality Control

To ensure the precise alignment of structural and diffusion MRI images a detailed visual quality control assessment was performed for each subject, following the guidelines provided in the ExploreDTI manual (8). This essential process involved overlaying the structural image onto the diffusion first eigenvector-fractional anisotropy (FEFA) map, as depicted in eFigure 1. The FEFA map was specifically selected for its ability to accentuate the directional orientation of diffusion fibres, enhancing the visibility of any potential misalignments between the structural and diffusion MRI images. Particular attention was paid to the alignment of the cerebrospinal fluid, pia layer, and major white matter tracts, ensuring their accurate co-registration. This meticulous cross-referencing was vital for precise alignment verification between the structural and diffusion MRI images.

Following visual inspection, 338 scans were found suitable for analysis, with a total of 22 scans removed; 17 of these were excluded due to unsuccessful connectome generation that could not be rectified, while an additional 5 were eliminated due to structural and diffusion image alignment discrepancies.

**eFigure 1.** An example of an appropriate structural and diffusion image alignments

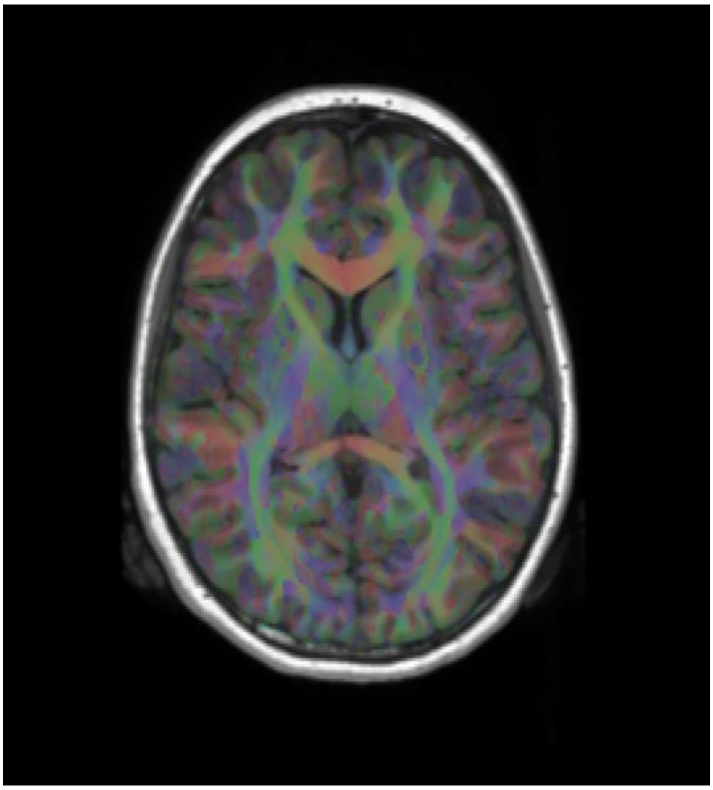

**eFigure 1.** This figure shows an example of an accurate alignment of a subject's structural and diffusion MRI images. As per the visual quality control strategy, subject-specific diffusion MRI FEFA and structural MRI images were overlaid and a visual inspection to confirm alignment accuracy.

#### 3. White matter tract extraction protocol

##### 3.1. Cingulum Bundles

The cingulum bundle is a prominent white matter tract interconnecting frontal, parietal, and medial temporal regions, as well as linking subcortical nuclei to the cingulate gyrus (9). The cingulum bundle is a key limbic system white matter tract, interconnecting the amygdala, hippocampus, cingulate gyrus as well as other regions (10). The 'standard cingulum' was reconstructed bilaterally using the protocol outlined by Jones and colleagues (11). To locate the rostral-caudal midpoint of the body of the corpus callosum, the midpoint between the posterior part of the curve of the genu (i.e., its most posterior part at flexion and the front of the splenium (i.e., its most anterior part at flexure) was identified. Two "AND" ROI – the term "AND" indicates that fibres must pass through this ROI – drawn five sections anterior and posterior of the identified rostral-caudal midpoint of the body of the corpus callosum. These two "AND" ROIs were separated by approximately 18mm in the anterior-posterior plane. All streamlines that passed through both "AND" ROIs were retained as 'cingulum bundle' pathways. Additional, "NOT" ROIs – the term "NOT" indicates that any fibres passing through this ROI are excluded – were used to exclude tracts that were inconsistent with known projections of the cingulum bundle. This procedure was repeated to extract the cingulum bundles in both hemispheres. A complete reconstruction of the bilateral cingulum bundles is provided in Figure 1 in the main manuscript.

##### 3.2. Fornix

The fornix is a key limbic system white matter bundle that interconnects the medial temporal lobe to the mammillary bodies and hypothalamus (10). The ROIs were manually drawn for each individual subject using landmark techniques that have been shown to be highly reproducible (12). A seed ROI – a term "SEED" indicates that fibres either originated or terminated in the ROI – was placed where the anterior pillars enter the fornix body (coronal plane). Using the mid-sagittal plane as a guide, an "AND" ROI was drawn in the axial plane encompassing both the crus fornici at the lower part of the splenium of the corpus callosum. Finally, two "NOT" ROIs were drawn, 1) rostral to the anterior fornix pillars and caudal to the crus fornici (coronal slice), 2) through the corpus callosum and the upper pons (axial slice) to exclude streamlines from the corpus callosum and corticospinal tract. Once all ROIs were placed the fornix was reconstructed. This procedure was repeated to extract the fornix in both hemispheres. Figure 1 in the main manuscript displays a complete reconstruction of the bilateral fornix.

##### 3.3. Mammillothalamic Tracts

The mammillothalamic tracts run inferior-superior from the mammillary bodies to the anterior thalamic nuclei and are the major connections of the anterior thalamic nuclei with the hypothalamic nuclei (13). The multiple ROI approach to extracting the mammillothalamic tracts was described by Kamali and colleagues (2018) (14). Using a subject-specific FEFA diffusion image the first “AND” ROI was drawn to encompass the fibres coursing through the mammillary bodies (axial plane). The second “AND” ROI was placed to contain the streamlines projecting from the medial aspect of the anterior thalamic nuclei. The addition of a second “ROI” excluded possible contamination from the stria terminalis and fornix. This procedure was repeated to extract the mammillothalamic projections in both hemispheres. An example of a reconstructed bilateral mammillothalamic projections is provided in Figure 1 in the main manuscript.

##### 3.4. Anterior Thalamic Projections

The anterior thalamic nuclei receives incoming fibres from the mammillothalamic tract and fornix and send outgoing projections via the anterior thalamic pathways to the orbitofrontal and anterior cingulate cortex (10). These anterior thalamic projections traverse through the anterior limb of the internal capsule, which is a dense bundle of white matter fibres that connects the thalamus to the cerebral cortex (10). A multiple ROI approach was used to extract the anterior thalamic projections (15). The first “AND” ROI was drawn that defines the anterior limb of the internal capsule (coronal plane), The second “AND” ROI was drawn, and the entire thalamus was delineated (coronal slice). A “NOT” ROI passing through the midbrain, the corticospinal tract and the corticopontine tract (axial slice) were set to remove anatomically implausible fibres. This procedure was repeated to extract the anterior thalamic projections in both hemispheres. Figure 1 in the main manuscript shows reconstructed bilateral anterior thalamic projections.

##### 3.5. Uncinate Fasciculus

The uncinate fasciculus interconnects the anterior region of the temporal lobe with the orbital and polar frontal cortex. The white matter fibres of the uncinate fasciculus originate from the temporal pole, uncus, parahippocampal gyrus, and amygdala. Projecting in a U-shaped turn, the fibres enter the floor of the external capsule (9). Two “AND” ROIs were placed as follows: 1) at the hippocampus-amygdala region, which was drawn in the temporal lobe at the junction with the anterior part of the temporal stem (axial slice), and 2) at the anterior cingulate area was drawn in the medial frontal lobe (coronal slice) (16). The two “AND” ROIs were drawn to cover a generous area to capture all possible streamlines that pass through both ROIs. This procedure was repeated to extract the uncinate fasciculi in both hemispheres. A complete reconstruction of the bilateral uncinate fasciculus is displayed in Figure 1 in the main manuscript.

#### 4. Statistical Analysis

##### 4.1. Mixed Model: Top-Down Model Selection Procedures

Primary statistical analyses were performed using the R software package (version 4.1.1) (17). To measure between-group differences in limbic system white matter, linear mixed-effects modelling (LMM) was performed using the *lme4* package in R (version 1.1-27.1) (18). A LMM is a versatile statistical approach that combines both fixed and random effects to analyse complex unbalanced data structures, such as those found in longitudinal studies (19). Fixed effects in an LMM represent the explanatory variables that are assumed to hold uniformly across the study such as sex and diagnostic group. These effects quantify the general relationships between these variables and the outcome of interest (DV), providing insights into broader patterns or tends within the data. Random effects, on the other hand, account for individual variations by allowing each subject to have their unique baseline (intercept) and rate of change over time (slope), thus capturing the inherent variability across subjects in the data.

An essential procedure of LMM is data-driven model selection, in which the goal is to identify a parsimonious model (i.e., high goodness of fit using as few explanatory variables as possible) to reduce the risk of a Type 1 error (20). An established top-down LMM model selection was used to select the optimal model for each structure of the limbic system (20). The details of the LMM models tested in this study are presented in eTable 1. IQ was not included as a covariate in any of the tested models as the inclusion of IQ is deemed inappropriate for neurodevelopmental disorders involving cognitive deficits such as ADHD as it can lead to overcorrected and spurious findings (21).

A top-down approach to model selection starts with the most complex model, which includes all random and fixed effects. These effects are then systematically removed in a backward fashion using a combination of fit statistics –Akaike Information Criterion (AIC) and Log-Likelihood Ratio test (LRT) – to identify the optimal model for each limbic system structure. The more complex model was selected if it provided a significantly better model fit compared to the simpler model (as indicated by having both an AIC > 5 and LRT p-value <0.05) (22-24). AIC and likelihood ratio tests were conducted by the default 'stats' package in R (v1.1-27.1) (17).

Given prior work suggesting non-linear relationships between brain structure and age (25, 26), linear and quadratic models were used to explore regional volumes were optimally fitted with quadratic function (age + I[$age^2$]) rather than linear (age) across the three time points of this study (Linear Model vs. Quadratic Model in Table ). The quadratic model was included if it significantly increased the model fit and the model's boundary was not singular.

Next, random effects were identified by comparing the fit of the models with and without the random effect of slope (RX1a vs. RX1b). The random effect of intercept and slope (RFX 1b) was included if it significantly increased model fit compared to the simpler model (RX1a). The fixed effects of interest (i.e., Diagnosis and Diagnosis-by-Age interaction) were identified by comparing the model fit of the null model (a model that only contained the covariate fixed effects: age, sex, age at baseline and intracranial volume [for connectome analysis]) against both fixed-effects models (FFX1/FFX2 vs. Null). If a fixed-effects model significantly increased fit compared to the null model, this indicated a significant effect on volume due to the fixed effects of interest. If both fixed effects models significantly increased model fit compared to the null model, but there was no difference in model fit between the two fixed effects models (FFX1 vs. FFX2), the parsimonious model (FFX1) was selected, as including the interaction term was not statistically justified. When initially comparing model fit, Maximum Likelihood (ML) was used to allow for comparability with the model fit parameters AIC and BIC. However, once the "optimal model" was identified, this model was refitted using Restricted Maximum Likelihood (REML) to increase the accuracy of estimates of the final model parameters. The covariance structure was set to "unstructured", which is the default covariance structure in R software. This structure imposes no restrictions on the covariance parameters, thus allowing the inclusion of variations in the slopes. In all the optimal models, a robust two-stage False Discovery Rate (FDR) (27) procedure was employed to correct for multiple comparisons at q = 0.05. Two-stage FDR correction was conducted using the *MuToss* package (28) in R (version 4.4). In line with recommendations for generating effect sizes in longitudinal mixed-effects modelling studies (29), pseudo-standardised coefficients (β) were calculated using the *effectsize* package (30) in R (version 4.1.1). These coefficients standardise according to the levels of the predictor, taking into account both within-group and between-group variants, thus offering accurate and reliable measure of effect sizes in longitudinal mixed-effects modelling studies.

##### 4.2. Brain-behaviour analyses

Brain-behaviour analyses were performed to examine the relationship between limbic system volumes and ADHD symptoms (CAI and ARI scores) in children and adolescents with ADHD. These relationships were investigated using LMM via the lme4 package in R (version 1.1-27.1) (18). The chosen LMM model, illustrated as model FX3b in eTable 1, evaluated if changes in limbic system volumes over time varied based on ADHD symptom severity, incorporating an age-by-ADHD symptoms interaction term. This model also adjusted for covariates age, sex, and intracranial volume. To mitigate the effects of multiple comparisons, a two-stage FDR correction was applied using the MuToss package in R (version 4.1.1)(28).

##### 4.3. Sensitivity analyses

Sensitivity analyses were performed to assess the potential impact of confounding factors (case-control sex imbalance and ADHD medication status) on the primary statistical analyses. The first sensitivity analysis examined the potential confound caused by the case-control sex imbalance in the study sample. To investigate this, 100 LMM iterations were run using randomly selected proportionately sex-matched case-control samples. To ensure the sex-matching in the control group was unbiased, female controls scans were randomly excluded at each iteration using a script developed in R studio (v.4.1.1). Averages of all 100 iterations were then collected to compare the results of the sensitivity analysis to the primary analysis. The second sensitivity analysis explored the potential impact of medication use in the ADHD group. This was accomplished by conducting LMM analyses on the optimal models to compare limbic system volumes between individuals with ADHD who were taking medication and those who were not. The third sensitivity analysis explored the potential impact of internalizing and externalizing comorbidities on limbic system white matter metric. To examine this, LMM analyses of the optimal models were conducted with an additional covariate of co-occurring disorder status (binary coded to indicate the presence or absence of an internalizing/externalizing disorder at Wave 1 and/or Wave 3). Finally, to investigate the impact of head motion, a sensitivity analysis was conducted, incorporating mean frame-wise displacement as a covariate in the primary analysis optimal models.

**eTable 1.** Mixed models tested: limbic system white matter metrics in ADHD and controls.

| | |
|---|---|
| **Higher Order Models** | |
| Linear Model | WM metric ~ intercept + $d_i$ + $b_1$(sex) + $b_2$(age) + $b_3$(age at baseline) + $b_4$(group*age) + $b_5$(ICV)* |
| Quadratic Model | WM metric ~ intercept + $d_i$ + $b_1$(sex) + $b_2$(age) + b3($age^2$) + $b_4$(age at baseline) + b5(group*age) + $b_6$(ICV)* |
| **Fixed Effects Models** | |
| Linear Null Model | WM metric ~ intercept + $d_i$ + $b_1$(sex) + $b_2$(age) + $b_3$(age at baseline) + $b_5$(ICV)* |
| Quadratic Null Model | WM metric ~ intercept + $d_i$ + $b_1$(sex) + $b_2$(age) + b3($age^2$) + $b_4$(age at baseline) + $b_6$(ICV)* |
| Linear FX simple Model | WM metric ~ intercept + $d_i$ + $b_1$(sex) + $b_2$(age) + $b_3$(age at baseline) + $b_4$(group) + $b_5$(ICV)* |
| Quadratic FX simple Model | WM metric ~ intercept + $d_i$ + $b_1$(sex) + $b_2$(age) + b3($age^2$) + $b_4$(age at baseline) + $b_5$(group) + $b_6$(ICV)* |
| Linear FX complex Model | WM metric ~ intercept + $d_i$ + $b_1$(sex) + $b_2$(age) + $b_3$(age at baseline) + $b_4$(group*age) + $b_5$(ICV)* |
| Quadratic FX complex Model | WM metric ~ intercept + $d_i$ + $b_1$(sex) + $b_2$(age) + b3($age^2$) + $b_4$(age at baseline) + b5(group*age) + $b_6$(ICV)* |
| Linear Brain-Behaviour Model | WM metric ~ intercept + $d_i$ + $b_1$(sex) + $b_2$(age) + b3($age^2$) + $b_4$(age at baseline) + b5(ADHD symptom score*age) + $b_6$(ICV)* |
| Quadratic Brain-Behaviour Model | WM metric ~ intercept + $d_i$ + $b_1$(sex) + $b_2$(age) + b3($age^2$) + $b_4$(age at baseline) + b5(ADHD symptom score *age) + $b_6$(ICV)* |

**eTable 1.** WM = white matter, RX = random effects, FX = fixed effects, ROI = regions of interest, ICV = intracranial volume, age = participant age from baseline (in months). To increase interpretability, the variables ICV and age at baseline were mean-centred.

### 5. Results

#### 5.1. Medication use during MRI scans

**eTable 2.** Medication use during diffusion MRI scans.

| Type of medication | Name of medication | Number of scans |
|---|---|---|
| Stimulants | | |
| | *Ritalin (Methylphenidate)* | 33 |
| | *Concerta (Extended-release Methylphenidate)* | 26 |
| | *Dexamphetamine* | 1 |
| | *Vyvanse (Lisdexamfetamine)* | 10 |
| Non-stimulants | | |
| | *Strattera (Atomoxetine)* | 5 |
| | *Guanfacine (Intuniv, Tenex)* | 2 |
| | *Catapres (Clonidine)* | 14 |
| Antidepressants | | |
| | *Lovan (Fluoxetine)* | 11 |
| Antipsychotics | | |
| | *Risperdal (Risperidone)* | 8 |
| Sleep Aids | | |
| | *Melatonin* | 20 |

#### 5.2. Non-significant between group analyses

**eFigure 2.** Non-significant between-group differences in DKI metrics.

**Axial Kurtosis**

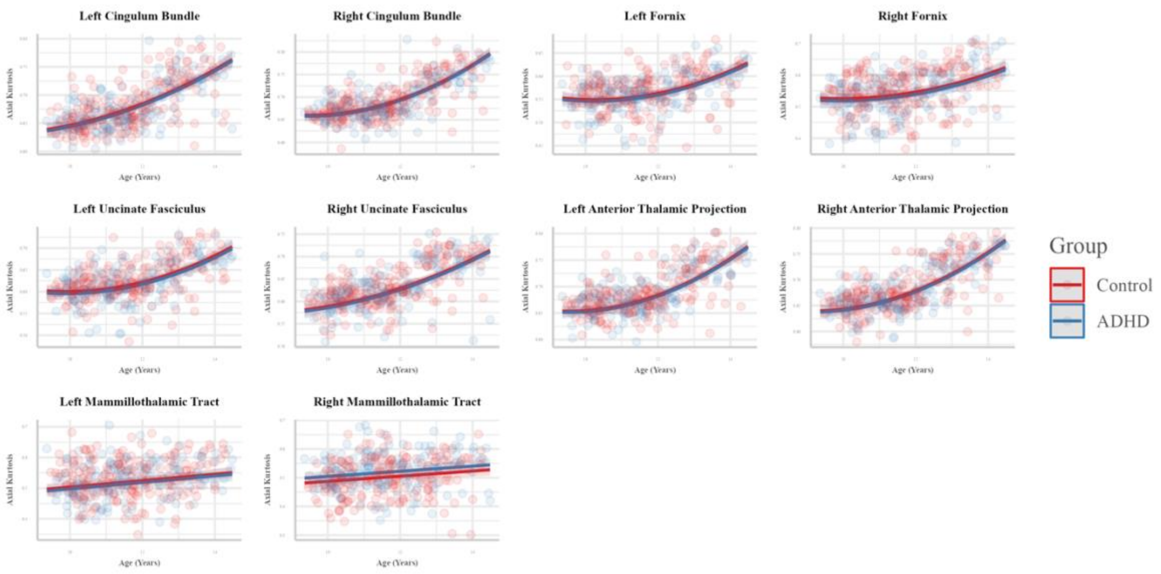

**Kurtosis Anisotropy**

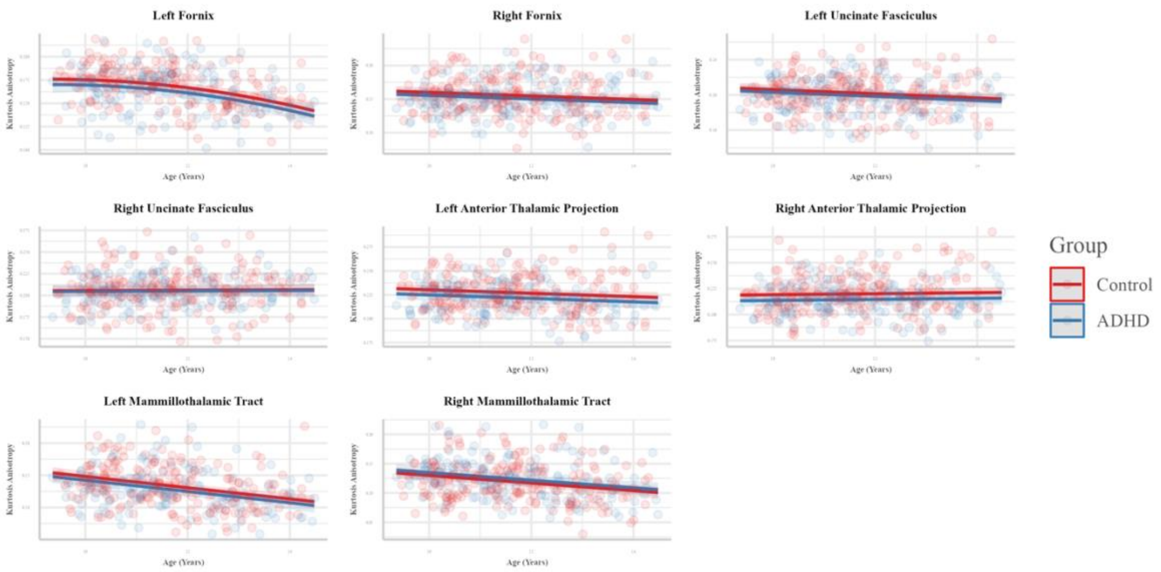

**Radial Kurtosis**

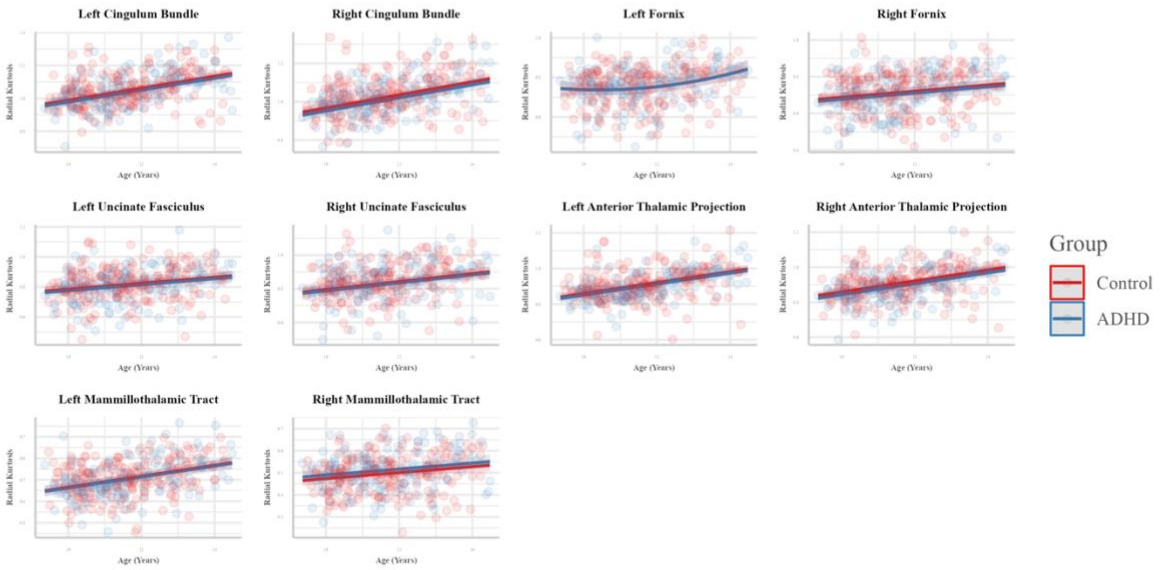

**Mean Kurtosis**

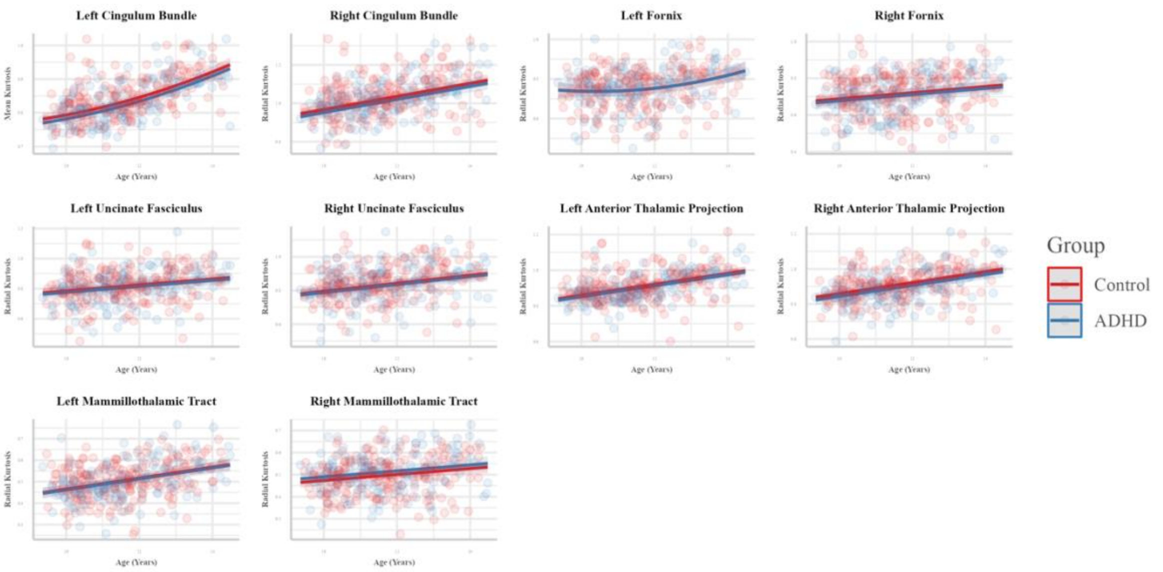

**Graph Theory Metics**

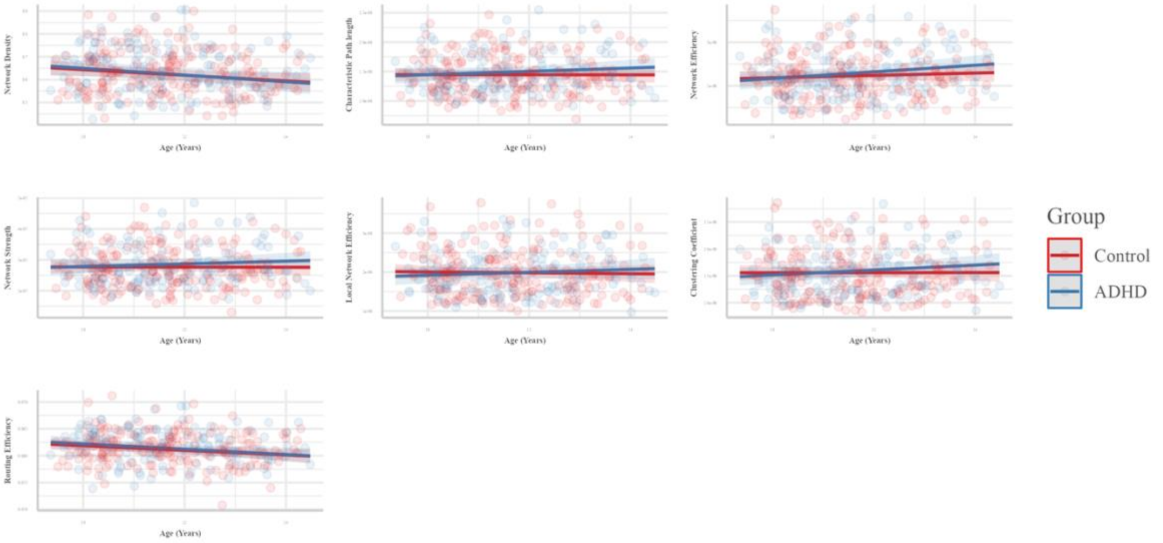

#### 5.3. Non-significant brain-behaviour analyses

**Figure 3.** Scatter plot of non-significant limbic system white matter DKI metrics by CAI scores in ADHD.

| Left Anterior Thalamic Projections | Right Anterior Thalamic Projections |
|---|---|

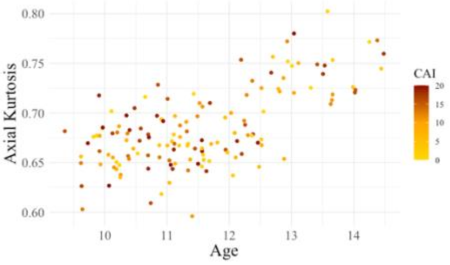

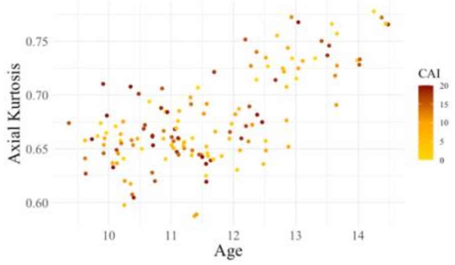

Left Fornix | Right Fornix

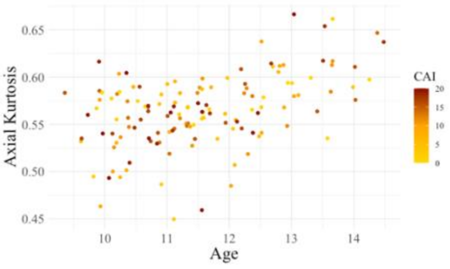

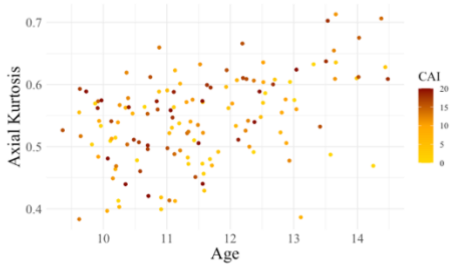

Left Uncinate Fasciculus | Right Uncinate Fasciculus

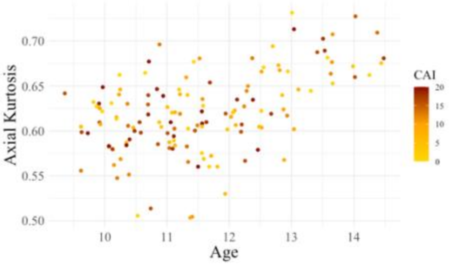

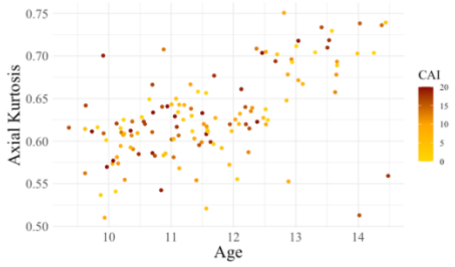

Left Mammillothalamic Tract | Right Mammillothalamic Tract

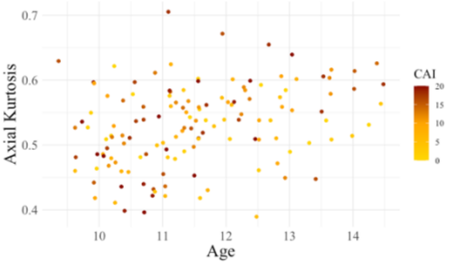

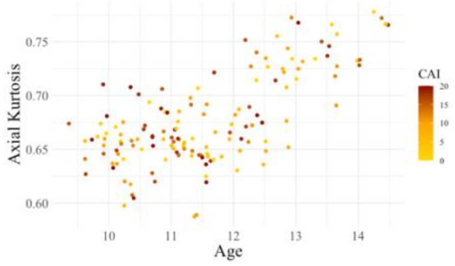

Left Cingulum Bundle | Right Cingulum Bundle

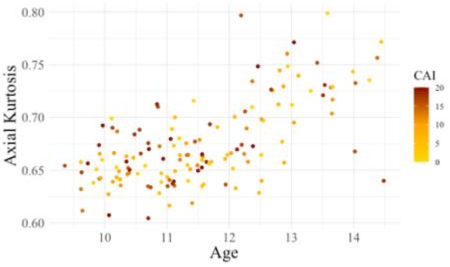

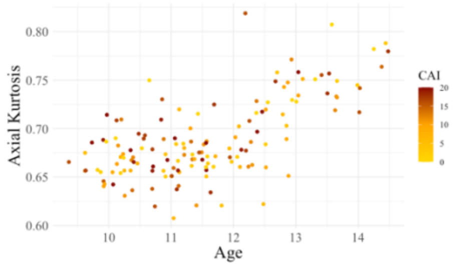

Left Anterior Thalamic Projections | Right Anterior Thalamic Projections

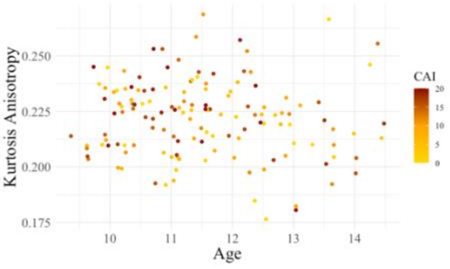

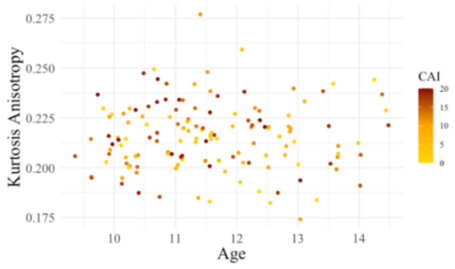

Left Fornix | Right Fornix

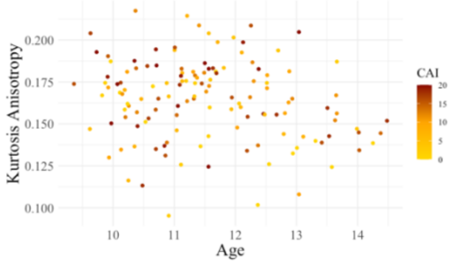

Left Uncinate Fasciculus

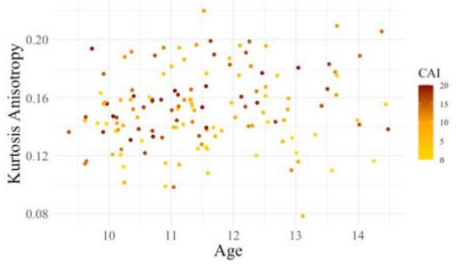

Right Uncinate Fasciculus

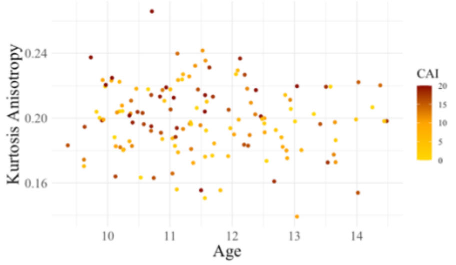

Left Mammillothalamic Tract

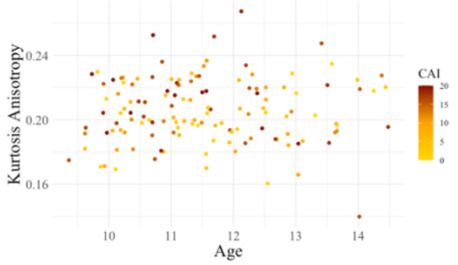

Right Mammillothalamic Tract

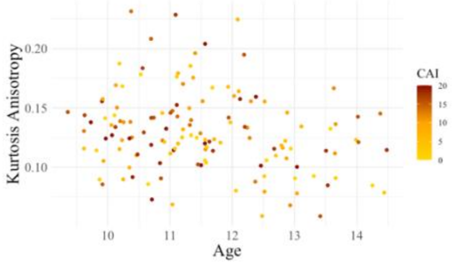

Left Cingulum Bundle

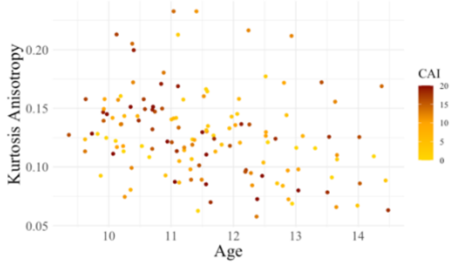

Right Cingulum Bundle

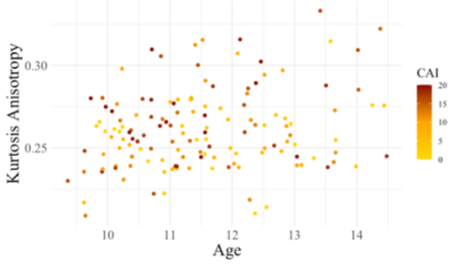

Left Anterior Thalamic Projections

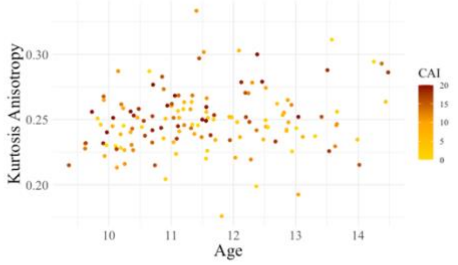

Right Anterior Thalamic Projections

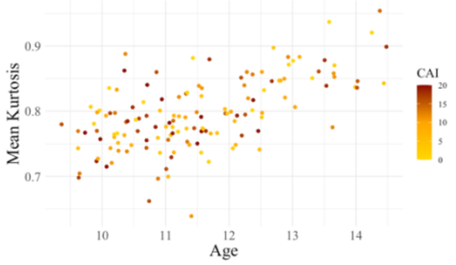

Left Fornix

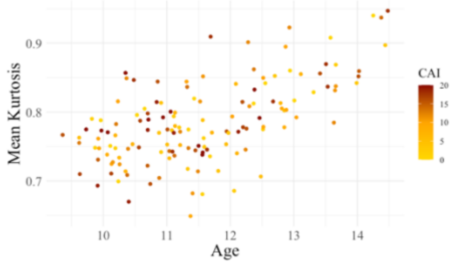

Right Fornix

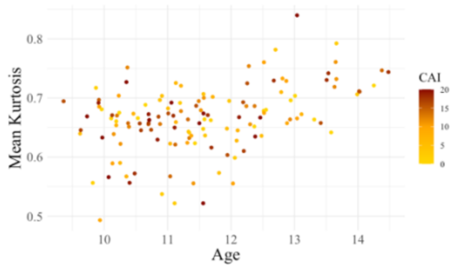

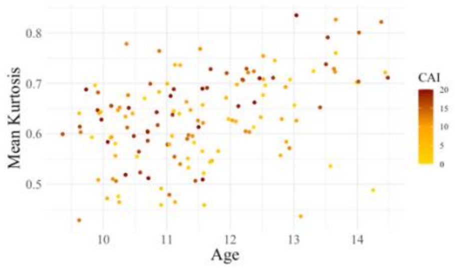

Left Uncinate Fasciculus | Right Uncinate Fasciculus

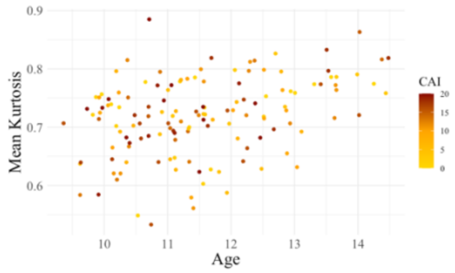

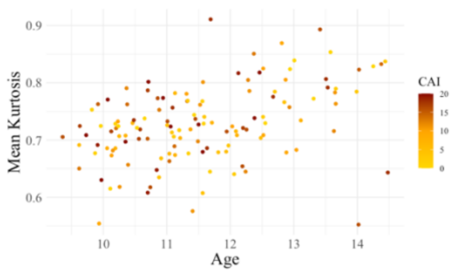

Left Mammillothalamic Tract | Right Mammillothalamic Tract

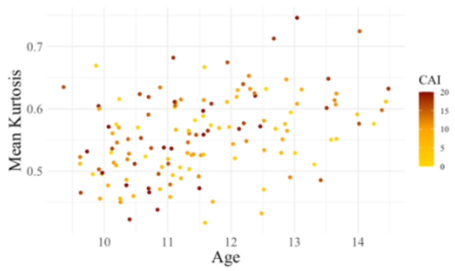

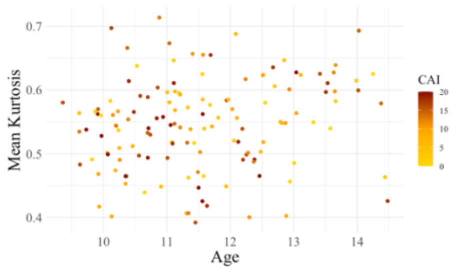

Left Cingulum Bundle | Right Cingulum Bundle

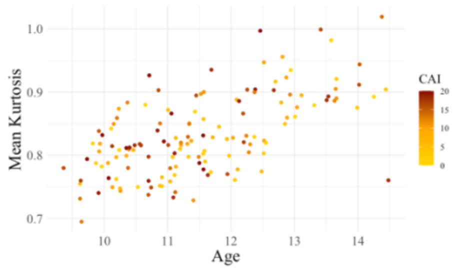

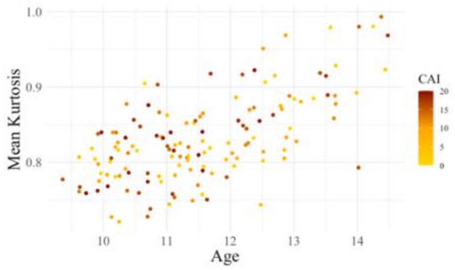

Left Anterior Thalamic Projections | Right Anterior Thalamic Projections

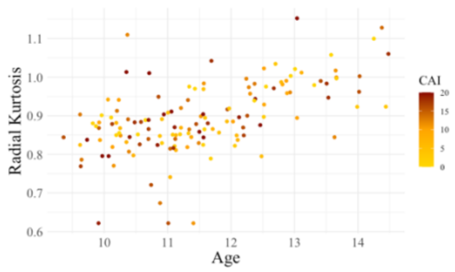

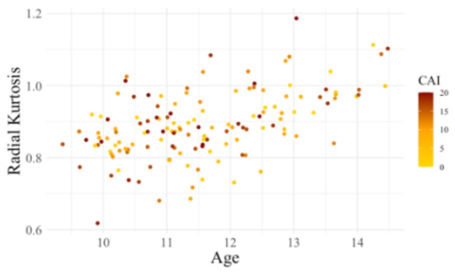

Left Fornix | Right Fornix

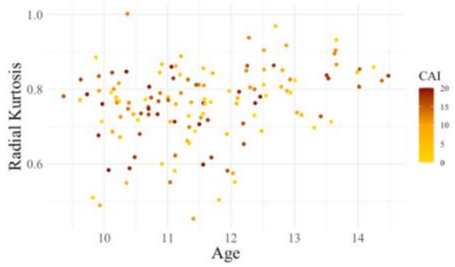

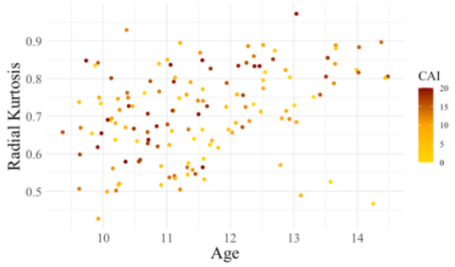

| Left Uncinate Fasciculus | Right Uncinate Fasciculus |
|---|---|

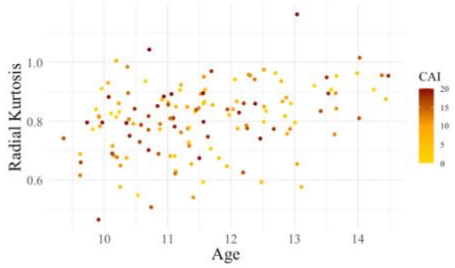

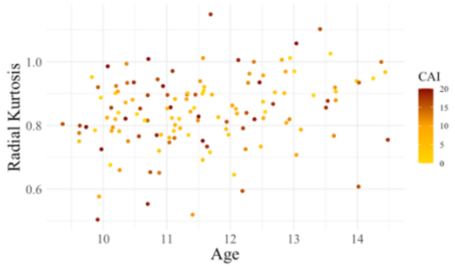

| Left Mammillothalamic Tract | Right Mammillothalamic Tract |
|---|---|

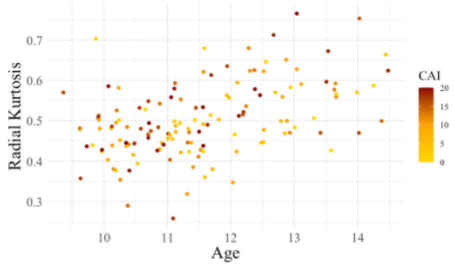

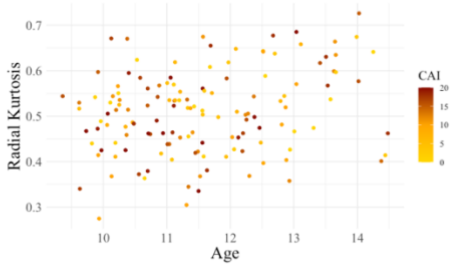

**eFigure 4.** Scatter plot of non-significant limbic system white matter DKI metrics by ARI scores in ADHD.

| Left Cingulum Bundle | Right Cingulum Bundle |
| --- | --- |

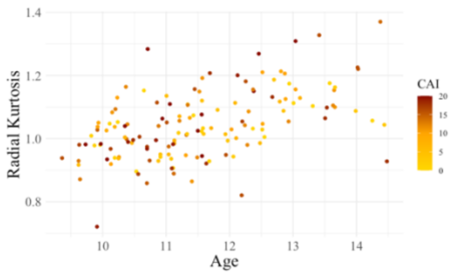

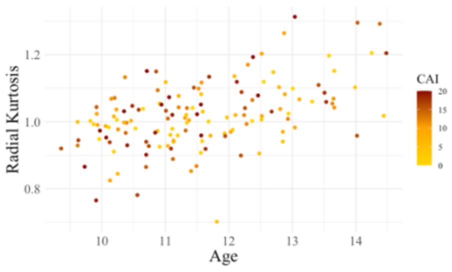

| Left Anterior Thalamic Projections | Right Anterior Thalamic Projections |
| --- | --- |

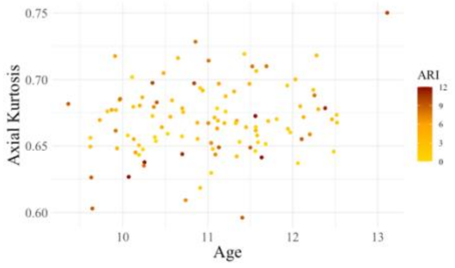

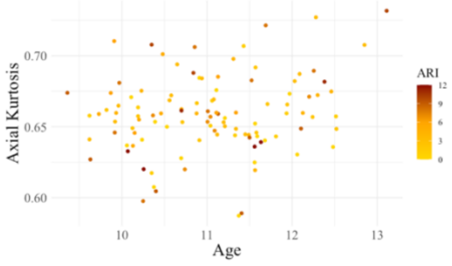

| Left Fornix | Right Fornix |
| --- | --- |

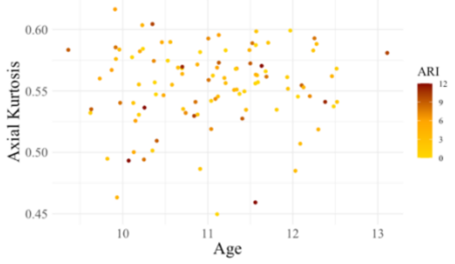

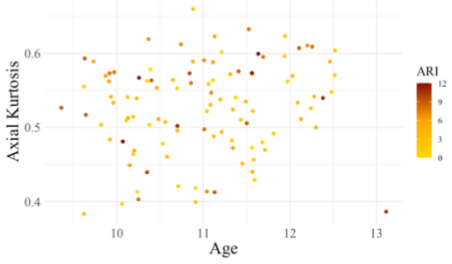

| Left Uncinate Fasciculus | Right Uncinate Fasciculus |
| --- | --- |

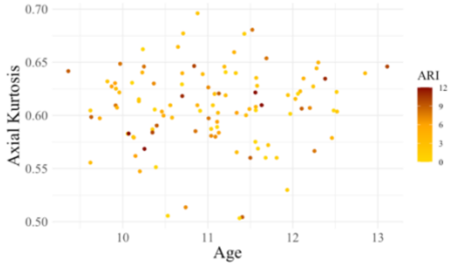

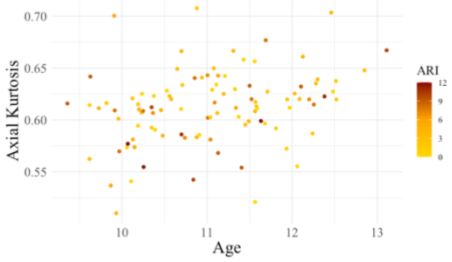

| Left Mammillothalamic Tract | Right Mammillothalamic Tract |
| --- | --- |

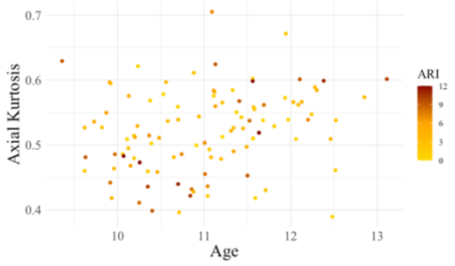

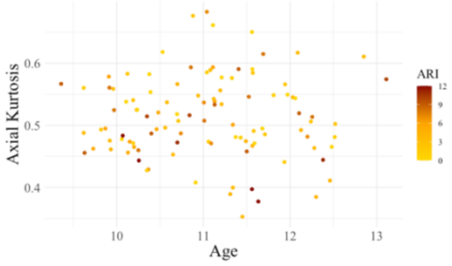

| Left Cingulum Bundle | Right Cingulum Bundle |
| --- | --- |

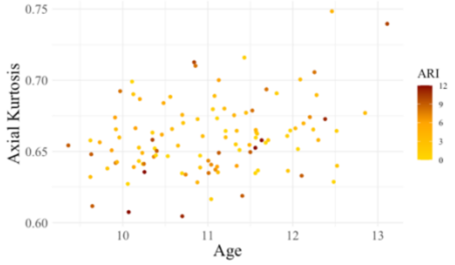

Left Anterior Thalamic Projections

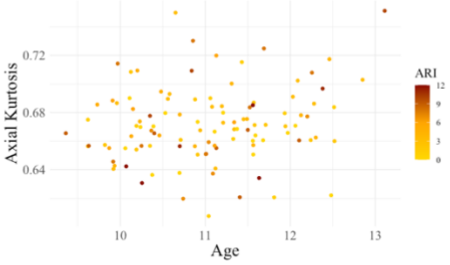

Right Anterior Thalamic Projections

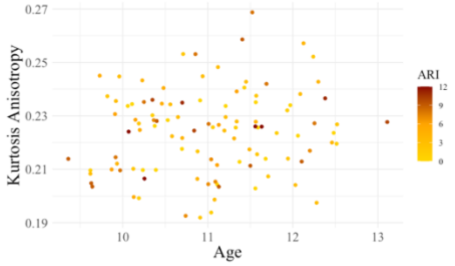

Left Fornix

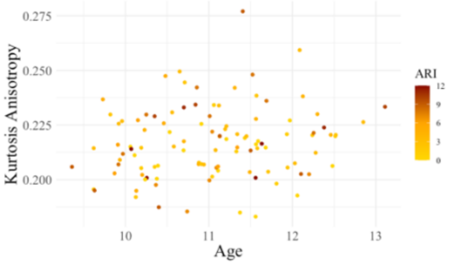

Right Fornix

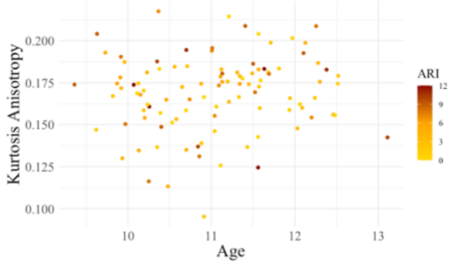

Left Uncinate Fasciculus

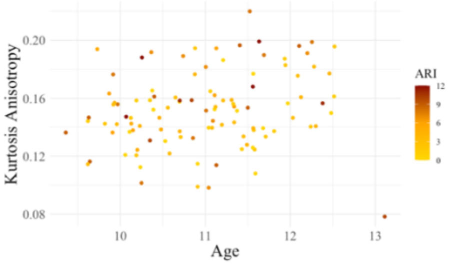

Right Uncinate Fasciculus

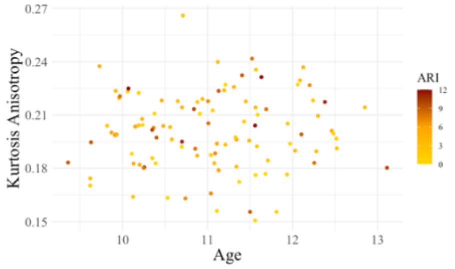

Left Mammillothalamic Tract

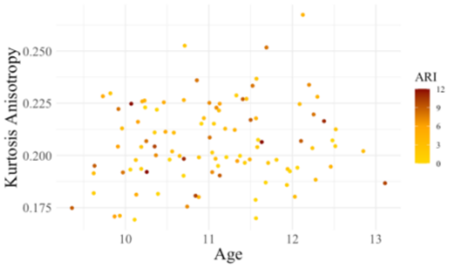

Right Mammillothalamic Tract

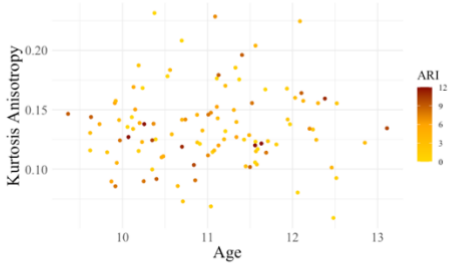

Left Cingulum Bundle

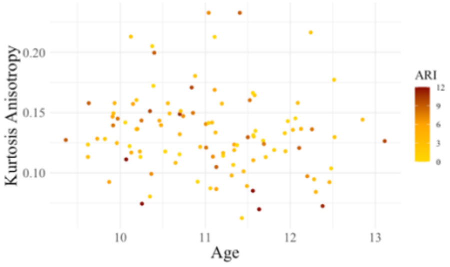

Right Cingulum Bundle

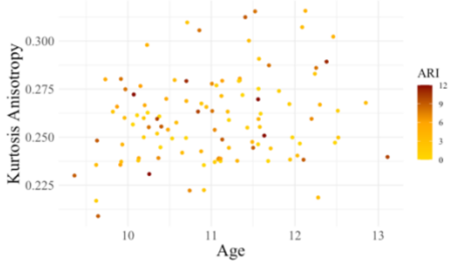

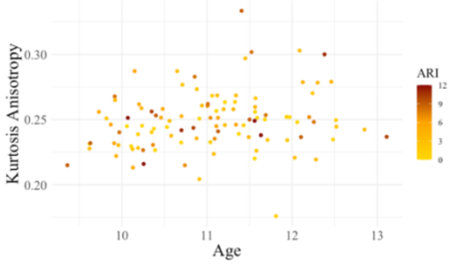

| Left Anterior Thalamic Projections | Right Anterior Thalamic Projections |
|---|---|

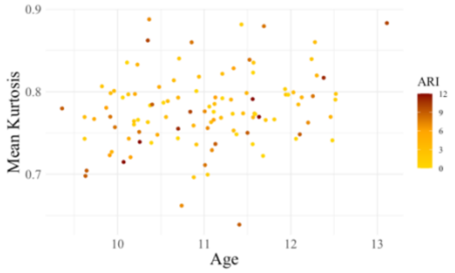

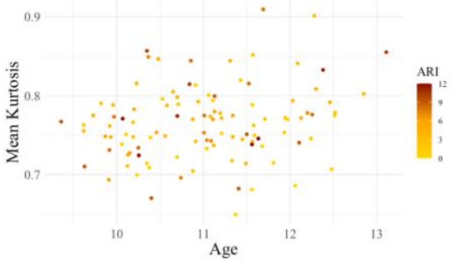

| Left Fornix | Right Fornix |
|---|---|

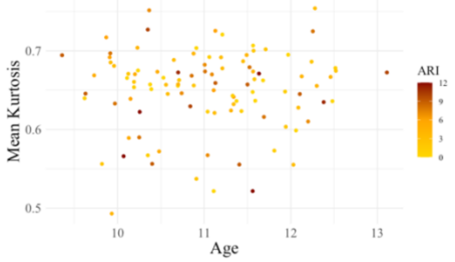

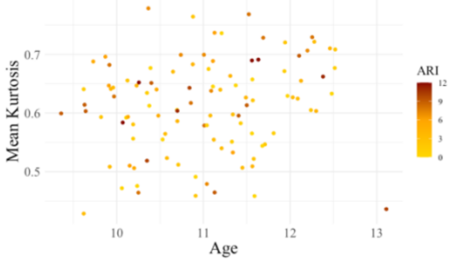

| Left Uncinate Fasciculus | Right Uncinate Fasciculus |
|---|---|

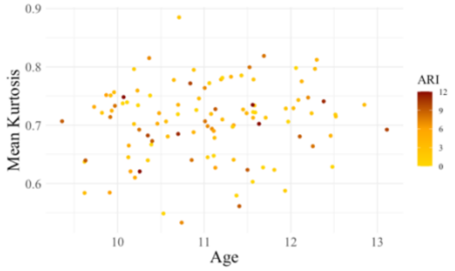

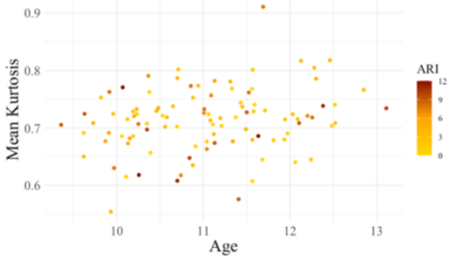

| Left Mammillothalamic Tract | Right Mammillothalamic Tract |
|---|---|

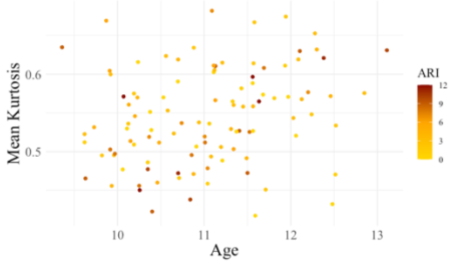

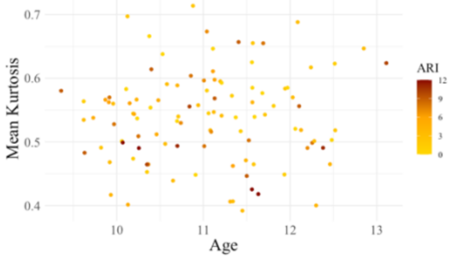

| Left Cingulum Bundle | Right Cingulum Bundle |
|---|---|

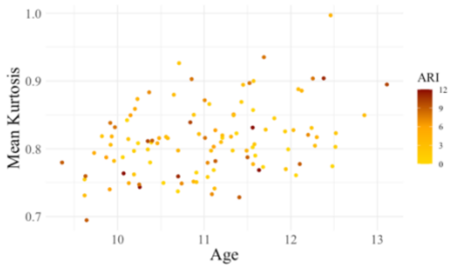

Left Anterior Thalamic Projections

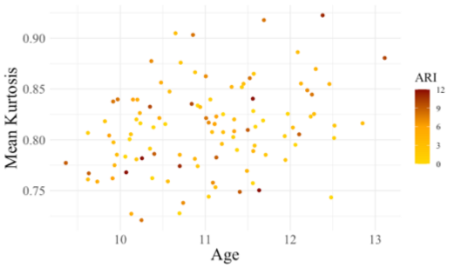

Right Anterior Thalamic Projections

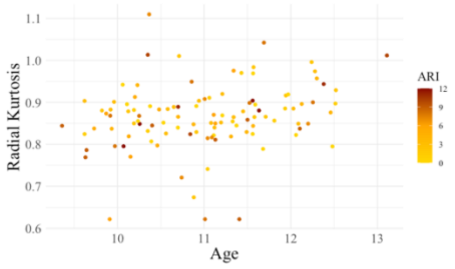

Left Fornix

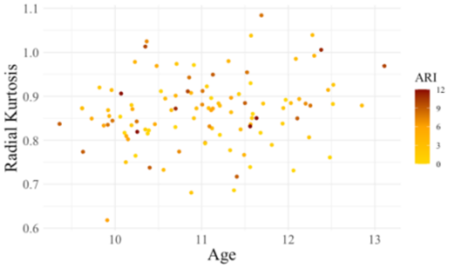

Right Fornix

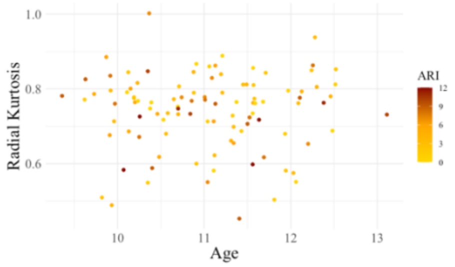

Left Uncinate Fasciculus

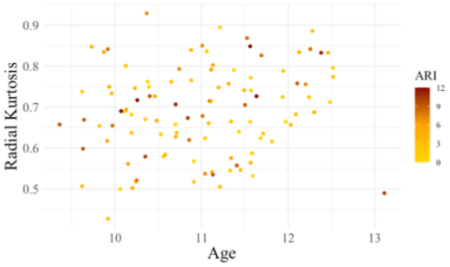

Right Uncinate Fasciculus

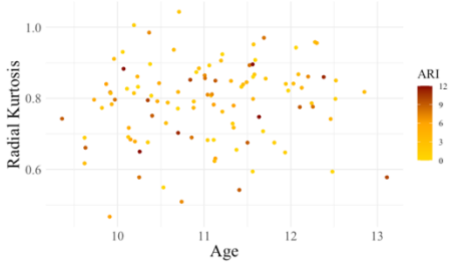

Left Mammillothalamic Tract

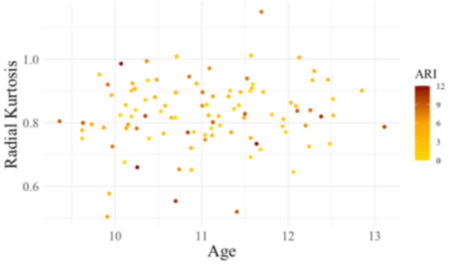

Right Mammillothalamic Tract

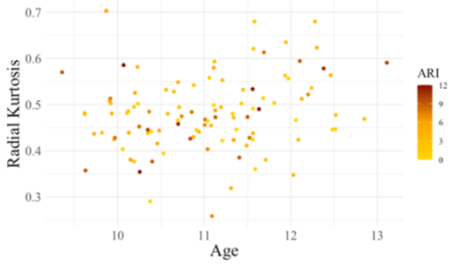

Left Cingulum Bundle

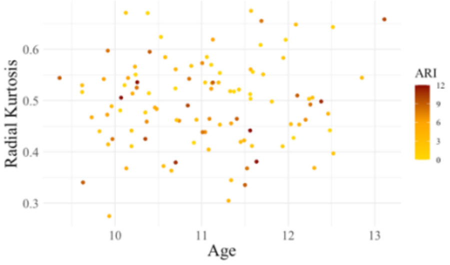

Right Cingulum Bundle

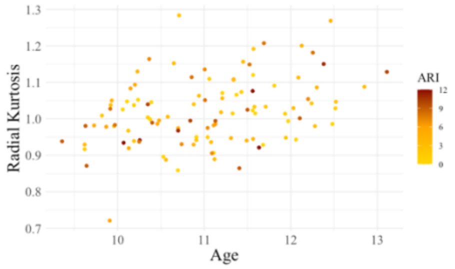

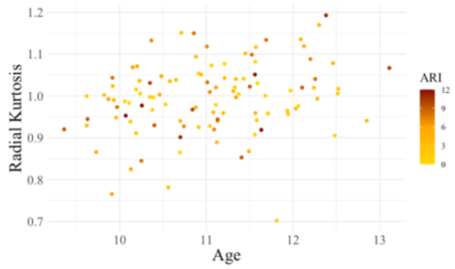

**eFigure 5.** Scatter plot of non-significant limbic system network measures by CAI scores in ADHD.

Network Efficiency

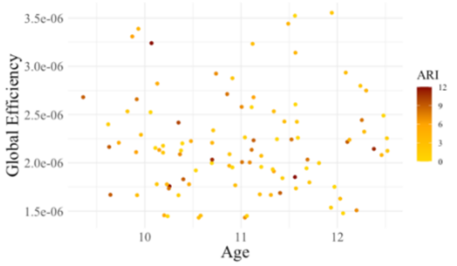

Characteristic Path Length

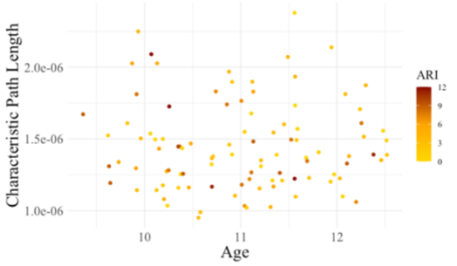

Clustering Coefficient

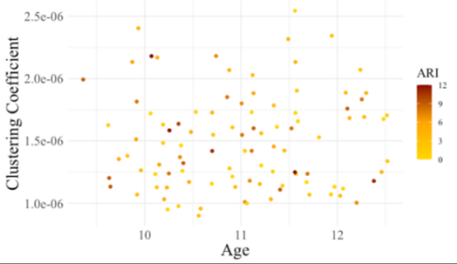

Network Strength

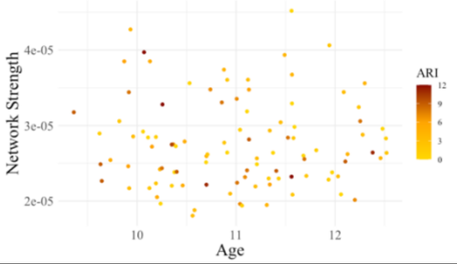

Local Efficiency

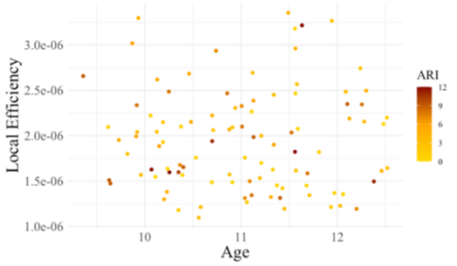

**eFigure 5.** This scatter plot exclusively features non-significant limbic system network measures (y-axis) alongside CAI scores (colour-coded) within the population. Each data point represents an individual in the study. The colour gradient used to represent CAI scores ranges from low (gold) to medium (orange) and high (dark red). Despite the absence of statistical significance, this visualisation sheds light on the relationship between non-significant limbic system network measures and CAI scores across the ADHD population.

**Figure 6.** Scatter plot of non-significant limbic system network measures by ARI scores in ADHD.

Network Efficiency

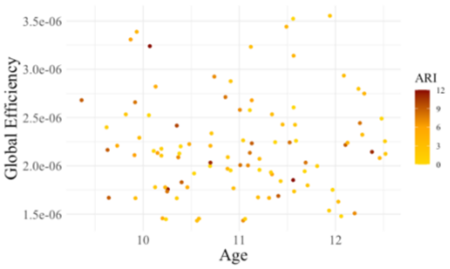

Characteristic Path Length

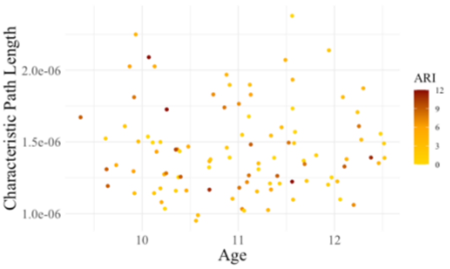

Clustering Coefficient

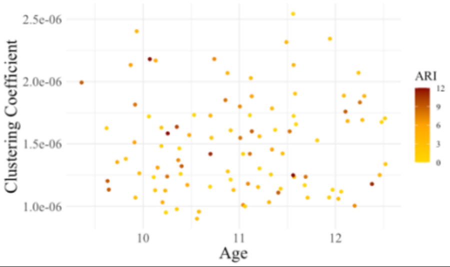

Network Strength

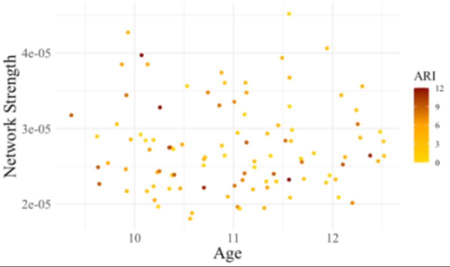

Network Density

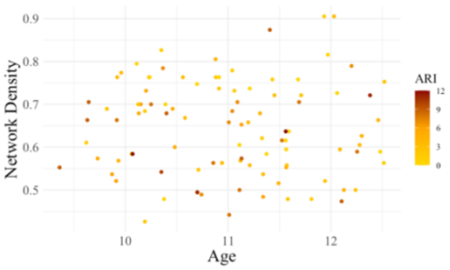

Routing Efficiency

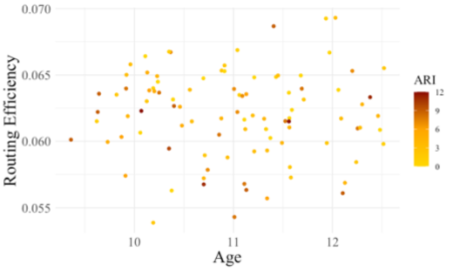

Local Efficiency

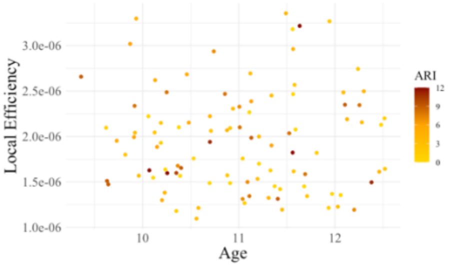

**eFigure 6.** This scatter plot exclusively features non-significant limbic system network measures (y-axis) alongside CAI scores (colour-coded) within the population. Each data point represents an individual in the study. The colour gradient used to represent CAI scores ranges from low (gold) to medium (orange) and high (dark red). Despite the absence of statistical significance, this visualisation sheds light on the relationship between non-significant.

#### 5.4. Results of sensitivity Analyses

*Microstructural DKI Between-Group Sensitivity Analysis*

The first sensitivity analysis, using the 100 random sex-matched linear mixed model analyses, reported an expected slight reduction in p-value significance, given the increase in standard errors (SE) related to the reduction in sample size (eTable 15). Importantly the beta values for the main effect of diagnosis were still within the SEs of the optimal models (Figure 7). Given the beta value and SEs, the sensitivity analysis demonstrated that the overall pattern of results in the primary analysis was most likely not confounded by a sex-ratio group imbalance in the study.

The second sensitivity analysis found there was no significant difference in white matter microstructural organization in the bilateral cingulum bundle among medicated ADHD and non-medicated ADHD children (eTables 16). The results of the third and fourth sensitivity analyses are presented in eTable 17 and 18, importantly in the sensitivity analyses the beta values for the effect of diagnosis remained within the SEs of the optimal models (see eFigures 8-9). Considering the beta value and SEs, this sensitivity analysis indicated that the main findings of the primary analysis were probably not confounded by the presence comorbidity or head motion. Overall, the sensitivity analyses demonstrated that the results of the primary analysis were most likely not biased by the sex ratio per group, medication status, comorbidity status or head motion.

*Macrostructural Graph Theory Brain-Behaviour Sensitivity Analyses*

The sensitivity analysis indicated that the medication status covariate did not drive the primary findings (eTable 19). Due to the collinearity between CAI scores and medication use ($r = 0.337$, $p = <0.001$), an expected reduction in p-value was reported when medication status was included as a covariate. The beta coefficients of the sensitivity analyses were largely unchanged and importantly remained within the standard errors of the main analyses. This suggests that the original relationship between network metric and CAI scores was relatively robust and not substantially affected by the inclusion of the covariate. The results from the second and third sensitivity analysis are presented in eTables 20 and 21. The sensitivity analyses revealed that the beta values associated with the main diagnostic effect were still within the SEs of the optimal models (eFigures 10 & 12). Based on the beta value and SEs, the sensitivity analysis suggested that the results of the exploratory analysis were likely not confounded by comorbidity status or head motion. Overall, the sensitivity analyses demonstrated that the results of the exploratory analysis were most likely not biased by medication status, comorbidity status or head motion.

**eTable 15.** Mean values of the effect of diagnosis in the 100 iterations of optimal mixed-effects models analyses of limbic system with sex sex-matched case-control samples.

| | Diagnosis | | | |
|---|---|---|---|---|
| | ***B*** | ***SE*** | ***T*** | ***p-value*** |
| **Left Cingulum Bundle (KA)** | -8.50e-03 | 3.397e-03 | -2.515 | 0.014 |
| **Right Cingulum Bundle (KA)** | -8.89e-03 | 3.452e-03 | -2.575 | 0.011 |

**eTable 16.** Results of medication status sensitivity analysis using optimal mixed-effects models: limbic system white matter KA in ADHD.

| | **Sex** | | **Months from baseline** | | **Age at baseline** | | **Medication Status** | |
|---|---|---|---|---|---|---|---|---|
| | ***B (SE)*** | ***t, p*** | ***B (SE)*** | ***t, p*** | ***B (SE)*** | ***t, p*** | ***B (SE)*** | ***t, p*** |
| **Left Cingulum Bundle (KA)** | -9.298e-04 (5.888e-03) | -0.158, 0.8750 | 2.454e-04 (9.812e-05) | 2.501, 0.014 | 9.517e-03 (4.983e-03) | 1.910, 0.060 | -1.651e-03 (4.642e-03) | -0.356 0.722 |
| **Right Cingulum Bundle (KA)** | 7.010e-03 (5.801e-03) | 1.208, 0.231 | 2.309e-04 (9.961e-05) | 2.317, 0.022 | 1.208e-02 (4.907e-03) | 2.462 , 0.016 | -4.265e-03 (4.580e-03) | -0.931, 0.353 |

**eTable 17.** Results of optimal mixed-effects models analyses with comorbidity status: limbic system white matter microstructure in ADHD and Controls.

| | **Sex** | | **Age at baseline** | | **Months from baseline** | | **Diagnosis** | | **Comorbidity** | |
|---|---|---|---|---|---|---|---|---|---|---|
| | ***B (SE)*** | ***t, p*** | ***B (SE)*** | ***t, p*** | ***B (SE)*** | ***t, p*** | ***B (SE)*** | ***t, p*** | ***B (SE)*** | ***t, p*** |
| **Left Cingulum Bundle (KA)** | 1.678e-04 (3.626e-03) | 0.046 0.963 | 6.959e-03 (3.664e-03) | 1.899 0.059 | 2.683e-04 (7.601e-05) | 3.530 <0.000 | -8.618e-03 (3.448e-03) | -2.500 0.013 | 1.765e-03 (2.629e-03) | 0.671 0.502 |
| **Right Cingulum Bundle (KA)** | 4.022e-03 (3.756e-03) | 1.071 0.285 | 1.070e-02 (3.801e-03) | 2.816, 0.005 | 3.949e-04 (8.315e-05) | -4.749 <0.000 | -8.679e-03 (3.601e-03) | -2.410 0.016 | 1.188e-03 (3.023e-03) | 0.393 0.694 |

**eTable 18.** Results of optimal mixed-effects models analyses with frame-wise displacement: limbic system white matter microstructure in ADHD and Controls.

| | **Sex** | | **Age at baseline** | | **Months from baseline** | | **Diagnosis** | | **FWD** | |
|---|---|---|---|---|---|---|---|---|---|---|
| | ***B (SE)*** | ***t, p*** | ***B (SE)*** | ***t, p*** | ***B (SE)*** | ***t, p*** | ***B (SE)*** | ***t, p*** | ***B (SE)*** | ***t, p*** |
| **Left Cingulum Bundle (KA)** | -1.103e-03 (3.526e-03) | -0313 0.754 | 6.937e-03 (3.562e-03) | 1.947 0.053 | 2.672e-04 (7.447e-05) | 3.588 <0.000 | -8.955e-03 (3.330e-03) | -2.689 0.007 | 8.961e-03 (6.506e-03) | 1.377 0.169 |
| **Right Cingulum Bundle (KA)** | 8.639e-04 (3.351e-03) | 0.258 0.796 | 1.208e-02 (3.486e-03) | 3.465 <0.000 | 4.130e-04 (6.058e-05) | 6.818 <0.000 | -6.431e-03 (3.166e-0) | -2.031 0.043 | 1.097e-03 (5.749e-03) | 0.191 0.848 |

**eFigure 7.** Beta values and SEs of sex-group sensitivity analysis.

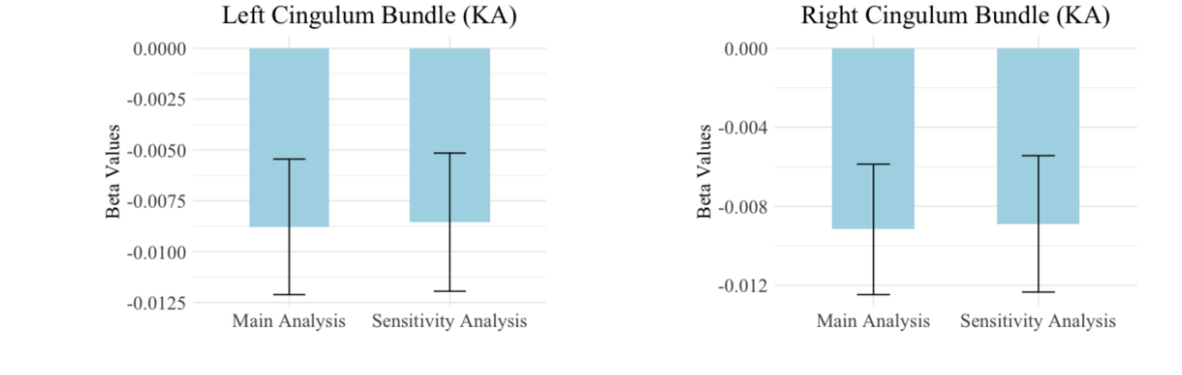

**eFigure 8.** Beta values and SEs of Kurtosis Anisotropy Comorbidity sensitivity analysis.

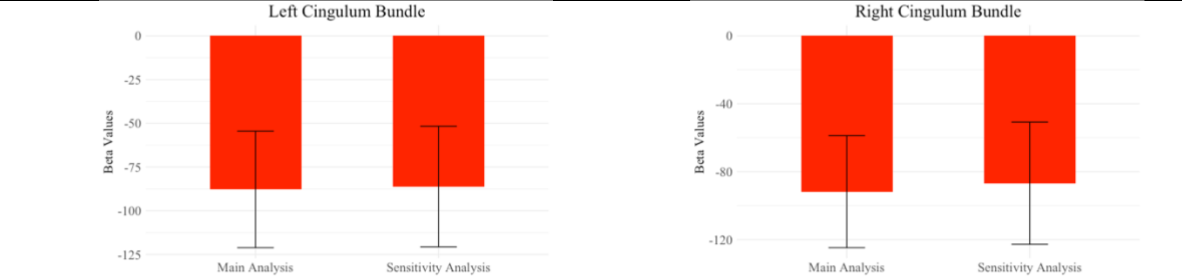

**eFigure 9.** Beta values and SEs of Kurtosis Anisotropy FWD sensitivity analysis.

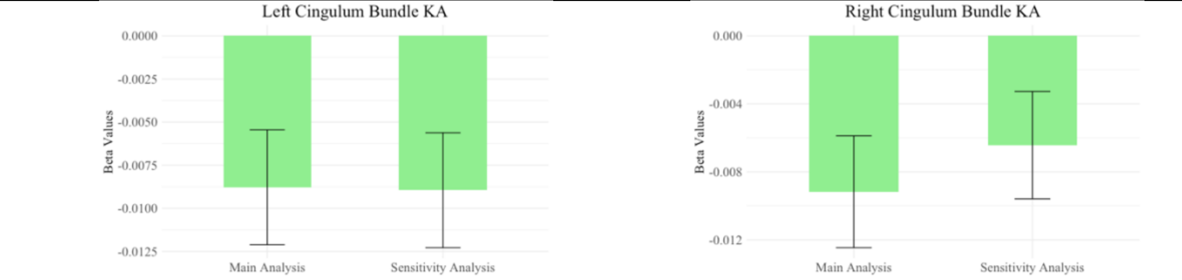

**eTable 19.** Results of mixed-effects models with medication status: limbic system network measures and CAI scores in ADHD.

| | ICV | | Sex | | Months from baseline | | Medication Status | | CAI | | Months from baseline * CAI | |
|---|---|---|---|---|---|---|---|---|---|---|---|---|
| | *B (SE)* | *t, p* | *B (SE)* | *t, p* | *B (SE)* | *t, p* | *B (SE)* | *t, p* | *B (SE)* | *t, p* | *B (SE)* | *t, p* |
| **Routing Efficiency** | -5.264e-10 (1.678e-09) | -0.314, 0.754 | 1.071e-03 (7.802e-04) | 1.373, 0.174 | 2.983e-09 (5.872e-09) | 0.508, 0.746 | -6.450e-05 (3.287e-05) | -1.962, 0.053 | -1.239e-04 (6.444e-05) | -1.923, 0.056 | 2.101e-06 (2.739e-06) | 0.767, 0.445 |
| **Network Density** | -1.584e-08 (5.684e-08) | -0.279, 0.781 | 3.486e-02 (2.574e-02) | 1.355, 0.180 | -2.138e-03 (1.204e-03) | -1.775, 0.079 | 2.053e-02 (2.487e-02) | 0.826, 0.411 | -4.586e-03 (2.224e-03) | -2.062, 0.041 | 9.823e-05 (9.878e-05) | 0.994, 0.322 |

**eTable 20.** Results of mixed-effects models analyses with comorbidity: limbic system network measures and CAI scores in ADHD.

| | ICV | | Sex | | Months from baseline | | Comorbidity | | CAI | | Months from baseline * CAI | |
|---|---|---|---|---|---|---|---|---|---|---|---|---|
| | *B (SE)* | *t, p* | *B (SE)* | *t, p* | *B (SE)* | *t, p* | *B (SE)* | *t, p* | *B (SE)* | *t, p* | *B (SE)* | *t, p* |
| **Routing Efficiency** | -1.330e-09 (1.709e-09) | -0.778 0.438 | 9.910e-04 (8.274e-04) | 1.198 0.235 | -6.924e-05 (3.330e-05) | -2.079 0.040 | -3.369e-04 (5.765e-04) | -0.584 0.560 | -1.644e-04 (6.296e-05) | -2.611 0.010 | 2.206e-06 (2.774e-06) | 0.795 0.428 |
| **Network Density** | -3.418e-08 (5.677e-08) | -0.602 0.548 | 3.177e-02 (2.648e-02) | 1.200, 0.234 | -2.165e-03 (1.221e-03) | -1.773, 0.079 | -8.249e-03 (2.003e-02) | -0.412, 0.681 | -4.931e-03 (2.145e-03) | -2.298, 0.023 | 9.913e-05 (1.001e-04) | 0.990, 0.324 |

**eTable 21.** Results of mixed-effects models analyses with FWD: limbic system network measures and CAI scores in ADHD.

| | **ICV** | | **Sex** | | **Months from baseline** | | **FWD** | | **CAI** | | **Months from baseline * CAI** | |
|---|---|---|---|---|---|---|---|---|---|---|---|---|
| | ***B (SE)*** | ***t, p*** | ***B (SE)*** | ***t, p*** | ***B (SE)*** | ***t, p*** | ***B (SE)*** | ***t, p*** | ***B (SE)*** | ***t, p*** | ***B (SE)*** | ***t, p*** |
| **Routing Efficiency** | -1.191e-09 (1.740e-09) | -0.684 0.495 | 8.991e-04 (8.267e-04) | 1.088 0.281 | -6.770e-05 (3.469e-05) | -1.952 0.054 | -5.236e-04 (1.961e-03) | -0.267 0.789 | -1.698e-04 (6.489e-05) | -2.617 0.010 | 2.353e-06 (2.887e-06) | 0.815 0.417 |
| **Network Density** | -2.246e-08 (5.742e-08) | -0.391 0.696 | 3.027e-02 (2.639e-02) | 1.147 0.256 | -2.107e-03 (1.254e-03) | -1.681 0.096 | -1.767e-02 (6.632e-02) | -0.266 0.790 | -5.214e-03 (2.192e-03) | -2.378 0.019 | 1.014e-04 (1.028e-04) | 0.987 0.326 |

**eFigure 10.** Beta values and SEs of medication sensitivity analyses.

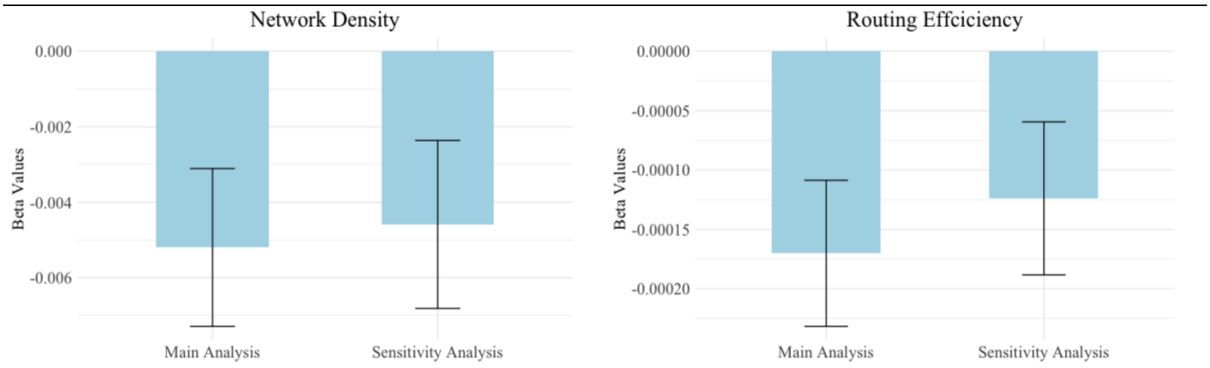

**eFigure 11.** Beta values and SEs of comorbidity sensitivity analyses.

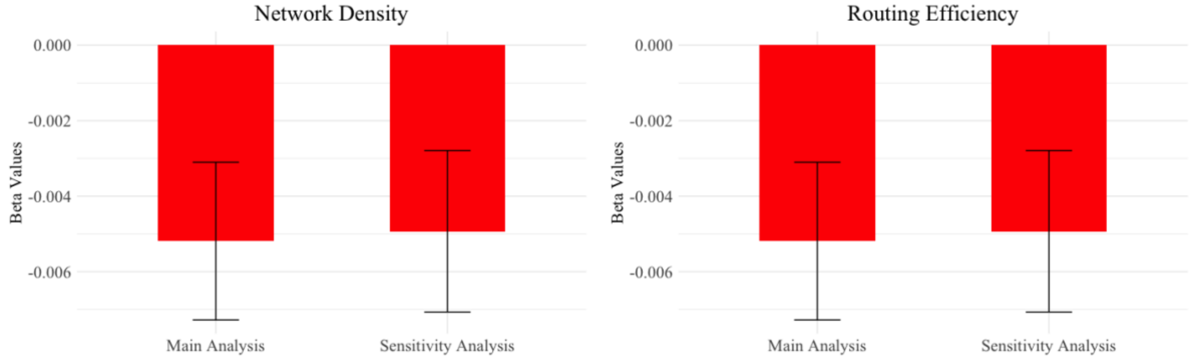

**eFigure 12.** Beta values and SEs of FWD sensitivity analyses.

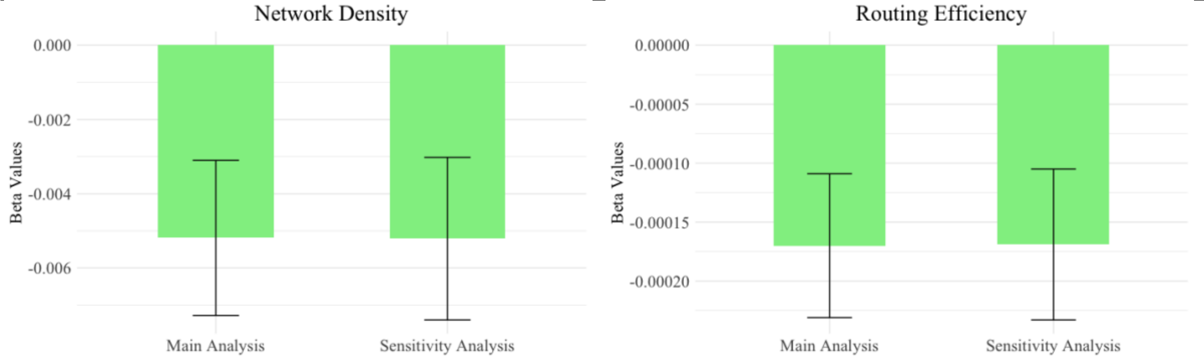
